# Pioneer factor IRF1 unlocks latent enhancers to rewire chromatin and immunometabolism in inflammatory macrophages

**DOI:** 10.64898/2026.02.27.708404

**Authors:** Jose-Mauricio Ayala, Rebecca Bellworthy, Mathieu Mancini, Ana Victoria Ibarra-Meneses, Christopher Fernandez-Prada, David Langlais

## Abstract

Macrophages undergo extensive chromatin and metabolic remodeling to mount effective inflammatory responses. Here, we identify Interferon Regulatory Factor 1 (IRF1) as a pioneer factor that orchestrates these processes during IFNγ-driven macrophage activation. Integrative ATAC-seq, ChIP-seq, Hi-ChIP, and nascent RNA-seq demonstrated that IRF1 rapidly engages closed chromatin, initiates enhancer remodeling, and drives removal of repressive histone marks followed by deposition of activating modifications. IRF1-established enhancers form long-range chromatin interactions with target promoters, activating transcriptional programs that control immune effector functions, chromatin regulation, and cellular metabolism. Notably, IRF1 coordinates the IFNγ-induced metabolic switch from oxidative phosphorylation to aerobic glycolysis by transcriptionally regulating key metabolic enzymes, with metabolite profiles consistent with increased glycolysis and pentose phosphate pathway engagement, while remodeling the tricarboxylic acid (TCA) cycle to support immunometabolic outputs. IRF1-deficient macrophages fail to execute this coordinated metabolic reprogramming. High-density IRF1 motif arrays promote enhanced chromatin occupancy and recruitment of the BRG1-containing chromatin remodeling SWI/SNF complex. Pharmacologic inhibition of SWI/SNF ATPase activity (SMARCA2/4) disrupts IRF1-dependent chromatin remodeling and gene induction. Moreover, IRF1-induced enhancer states persist after IFNγ withdrawal, establishing durable epigenetic memory. Together, these findings establish IRF1 as a central integrator of chromatin remodeling, transcriptional control, and metabolic adaptation, linking pioneer-factor activity to macrophage plasticity and innate immune memory.

## INTRODUCTION

The immune system depends on macrophages to coordinate pathogen clearance and inflammatory responses (Park et al., 2022). Alterations in chromatin status surrounding regulatory elements termed “enhancers” largely determine macrophage identity and activity by regulating transcriptional programs (Heinz et al., 2015; Lavin et al., 2014). The chromatin landscape of macrophages is primarily established during lineage commitment by lineage-determining transcription factors (LDTFs), such as PU.1 and C/EBPα, which create a permissive enhancer repertoire that is subsequently refined and activated by signal-dependent transcription factors (SDTFs) in response to developmental and environmental cues (Heinz et al., 2013; Mayran et al., 2019). In addition to pre-established enhancers, macrophages harbor a pool of “latent enhancers”, which are genomic regions lacking basal enhancer marks that can acquire chromatin accessibility and regulatory activity *de novo* upon specific inflammatory stimulation (Ostuni et al., 2013). An elite class of transcription factors (TF) known as pioneers, many of which belong to the LDTF category, can recognize their binding motifs within nucleosomal DNA in the context of compact heterochromatin (Ozden et al., 2023). Pioneer engagement initiates local chromatin relaxation and recruits chromatin remodeling complexes, thereby exposing regulatory elements to SDTF binding. SDTFs then exert regulatory activity by recruiting other transcription factors, chromatin-modifying proteins, and components of the transcriptional machinery (Ha et al., 2019).

IRF1 is a signal-dependent transcription factor with a crucial, non-redundant role in macrophage activation in response to some pro-inflammatory cues (Forero et al., 2019; Langlais et al., 2016; Roy et al., 2015; Tallam et al., 2016). Mice lacking IRF1 exhibit profoundly impaired interferon responses, particularly impacting type II interferon (IFNγ)-mediated immunity, making them highly susceptible to intracellular pathogens, including *Mycobacterium tuberculosis* and *M. bovis*, used in the BCG vaccine (Kano et al., 2008; Lohoff et al., 1997; Yamada et al., 2002). Similarly, humans with IRF1 deficiency develop severe, recurrent mycobacterial infections, as demonstrated by recent studies of children with inborn errors of immunity (Rosain et al., 2023). Early gene-specific studies demonstrated that IRF1 deficiency leads to reduced expression of critical chemokines and cytokines, including IL-12, IL-18, RANTES, CXCL9, CXCL10, and TNF-α, as well as impaired production of reactive oxygen species (ROS) (Chu et al., 2021; Forero et al., 2019; Kano et al., 2008; Liu et al., 2005; Maruyama et al., 2003; Roy et al., 2015; Siegmund et al., 2004; Yamada et al., 2002).

The advent of next-generation sequencing has shifted focus from isolated gene-centric analyses to genome-wide interrogation of transcription factor activity, including that of IRF1. Langlais *et al*. reveal extensive recruitment of IRF1 to chromatin in mouse bone marrow-derived macrophages (BMDMs) following IFNγ stimulation, highlighting its role in inducing histone H3K27 acetylation and activity across thousands of genomic sites (Langlais et al., 2016). RNA-seq further established IRF1 as essential for the transcriptional activation of hundreds of immune-related genes. Song *et al*. extended these findings using ATAC-seq to show that IRF1 influences chromatin accessibility at numerous interferon-stimulated gene (ISG) promoters (Song et al., 2021). Together, these studies position IRF1 as a central regulator of IFNγ-induced chromatin states, yet in contrast to PU. 1, which predefines chromatin states during myeloid development (Feng et al., 2008; Ha et al., 2019; Minderjahn et al., 2020), the role of IRF1 in actively reshaping macrophage chromatin during activation remains largely unexplored.

Recent work underscores the tight functional coupling between chromatin remodeling and metabolic reprogramming in macrophages (Arts, Novakovic, et al., 2016; Dong et al., 2020; Donohoe & Bultman, 2012; Netea et al., 2020; Noe & Mitchell, 2019). In the case of pro-inflammatory stimulation, macrophages undergo a metabolic shift from oxidative phosphorylation toward aerobic glycolysis, a transition that is essential for generating the energy and biosynthetic intermediates required to sustain pro-inflammatory responses and epigenomic changes (Mills et al., 2016; Palsson-McDermott et al., 2015; Tannahill et al., 2013; Wang et al., 2018). Consistent with this paradigm, blockade of glycolysis, either by glucose deprivation or pharmacological inhibition, disrupts IFNγ–driven macrophage activation, leading to reduced STAT1 phosphorylation, IL-1β, iNOS, and NO production (Wang et al., 2018). Despite this interdependence, the transcriptional and epigenetic mechanisms that link pro-inflammatory signaling to macrophage metabolic reprogramming remain poorly defined.

In this study, we investigate how IRF1 shapes epigenetic, transcriptional, and metabolic reprogramming in primary mouse macrophages in response to IFNγ stimulation. Using integrated multi-omics approaches, we show that IRF1 exhibits a spectrum of genome-wide binding behaviors, including strong binding at low-accessibility genomic regions, a property that underlies its pioneering activity. IRF1 binds closed heterochromatin to unlock latent enhancers, driving chromatin accessibility and establishing new regulatory elements that control key macrophage effector functions, including host defense, chromatin modifiers, and oxidative stress responses. In parallel, IRF1 engages pre-established enhancers at metabolic genes, coordinating transcriptional programs that promote glycolysis, pentose phosphate pathway (PPP) activity, and rewiring of the TCA cycle to support redox balance and immunometabolite production. In the absence of IRF1, macrophages fail to transition from oxidative phosphorylation to glycolysis, positioning IRF1 as a central mechanistic link between pro-inflammatory signaling and metabolic adaptation. Mechanistically, we demonstrate that IRF1 is required for recruitment of the BRG1-containing SWI/SNF chromatin remodeling complexes to these regulatory elements, and that SWI/SNF activity is necessary for full IRF1-dependent chromatin remodeling and gene induction, as demonstrated by pharmacological inhibition of BRG1. Finally, we show that IRF1-driven epigenetic remodeling and metabolic adaptations persist for days after IFNγ withdrawal, revealing a durable epigenetic state reminiscent of trained immunity. Together, these findings define IRF1 as a signal-dependent pioneer TF that links inflammatory signaling to stable chromatin and metabolic reprogramming in activated macrophages.

## RESULTS

Previous human and mouse studies have shown that loss of IRF1 renders macrophages unable to respond to some activation cues and clear intracellular pathogens, resulting in severe immunodeficiency. To better understand the mechanistic basis of IRF1’s essential role in macrophage activation, we applied a multi-omics approach to primary murine macrophages undergoing IFNγ-induced activation.

### IRF1 promotes chromatin accessibility at IFNγ-responsive regions in macrophages, independent of pioneer factor PU.1

To define how IRF1 regulates macrophage activation, we profiled genome-wide chromatin accessibility by ATAC-seq in wild-type (WT) and *Irf1-/-* (IRF1 KO) bone marrow-derived macrophages (BMDMs) across an extensive 0 to 48 h IFNγ stimulation time course (**Figure 1A**; **Supplementary Figure 1A**). Across all conditions, we identified a total of 140,489 accessible regions, of which 107,895 met high-confidence criteria (**Figure 1B**). Unsupervised hierarchical clustering of the ATAC-seq signal partitioned these sites into IFNγ-responsive (38,564) and IFNγ-unresponsive (69,331) fractions (**Figure 1B**). The unresponsive regions (64.3% of sites) showed stable accessibility in response to IFNγ and were independent of IRF1 (**Figure 1C**).

**Figure 1.**
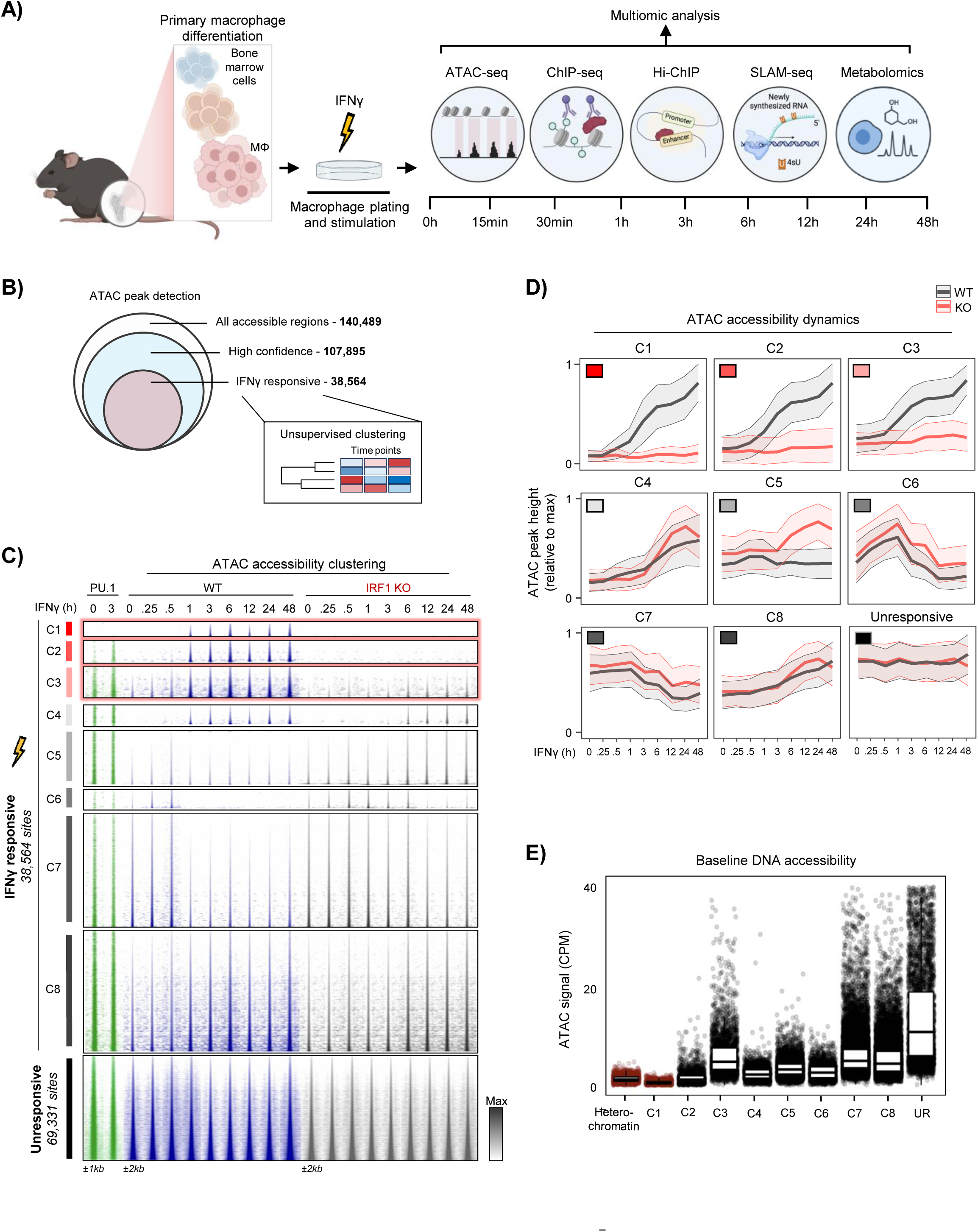
IRF1 drives accessibility at IFNγ-responsive chromatin in macrophages independent of myeloid pioneer PU.1. **(A)** Experimental design for multiomic assessment of WT and IRF1 KO bone marrow–derived macrophages (BMDMs) response to IFNγ stimulation. Time-resolved profiling by ATAC-seq, ChIP-seq, Hi-ChIP, SLAM-seq and metabolomics via GC/LC-MS is performed. **(B)** Venn diagram summarizing ATAC-seq–identified accessible chromatin regions, filtered for high-confidence peaks and IFNγ-responsiveness (n=38,564); this set is used for downstream clustering and differential analyses. **(C)** Heatmap of normalized ATAC-seq signal (rows = individual accessible site; columns = time points), grouped into eight clusters by k-means clustering. Clusters C1-C3 show IRF1-depedent increase in accessibility in response to IFNγ; highlighted in red. PU.1 ChIP-seq binding signal is also shown, with Cluster C1 lacking detectable PU.1 occupancy. **(D)** Ribbon plots of relative ATAC-seq peak height (each peak scaled to its maximum) over matched time points; lines indicate mean accessibility and shaded ribbons show ± SD for WT (black) and IRF1 KO (red). **(E)** Boxplots of normalized ATAC-seq counts in WT BMDMs at heterochromatin regions, and at clusters C1–C8 and unresponsive ATAC-seq sites; median with interquartile range are shown.

The remaining 35.7% of regions exhibited IFNγ-responsive chromatin, with a subset displaying IRF1 dependency (**Supplementary Figure 1B**). To resolve these dynamics, the IFNγ-responsive sites were subdivided into eight clusters (C1-C8) based on accessibility kinetics and dependency on IRF1 (**Figure 1C**; example of peaks for each cluster provided in **Supplementary Figure 1C**). Strikingly, clusters C1, C2, and C3 showed the strongest increase of chromatin accessibility in response to IFNγ, with pronounced reductions in *Irf1^-/-^* macrophages (**Figure 1D**). This dependency was most striking at C1 sites, which were fully dependent on IRF1 for accessibility gains. In contrast, accessibility gains at clusters C4 and C8 and losses at C6 and C7 were largely independent of IRF1. Cluster C5 accessibility remains low and stable in WT macrophages but interestingly increases in *Irf1^-/-^*upon IFNγ treatment (**Figure 1C-D**).

Because the macrophage epigenetic landscape is established by pioneer TFs, mainly PU.1, we integrated previously published PU.1 ChIP-seq datasets from resting and IFNγ-stimulated BMDMs. In line with its pioneer role, PU.1 occupancy was detected at most responsive and non-responsive regions, with the notable exception of cluster 1 sites (**Figure 1C**), which lacked PU.1 signal. Consistently, C1 regions displayed the lowest baseline accessibility signal across all clusters, with levels comparable to heterochromatin (**Figure 1E**). These results identify IRF1 as a dominant driver of IFNγ-induced chromatin accessibility, including at a subset of inaccessible sites at baseline and that are independent of the canonical pioneer factor PU.1.

### Footprinting and motif analyses reveal hierarchical IRF1-driven IFNγ responses

We performed motif enrichment and ATAC footprinting analyses to identify factors regulating chromatin accessibility in macrophages responding to IFNγ. Motif network visualization across ATAC clusters revealed a strong enrichment of IRF1 motifs in the IRF1-dependent clusters (**Figure 2A**). More than 50% of C1 sites contained a canonical IRF1 motif, with ∼20% harboring multiple IRF1 motifs (**Figure 2A**). Consistently, HOMER *de novo* motif analysis showed IRF-type motifs at the majority of C1 sites, whereas all other motifs were detected at frequencies of 1% or lower (**Supplementary Figure 2A**). C2 and C3 clusters displayed progressively lower IRF1 motif prevalence, with C2 characterized by a cooperative motif network involving PU.1, STAT1, and AP-1 yet still predominated by IRF1. C3 sites exhibited a more balanced contribution of IRF1, PU.1, and AP-1 (**Figure 2A**). By contrast, the unresponsive cluster was enriched for a diverse set of developmental and homeostatic TF motifs, including RUNX, PU.1, NFIL3, and AP-1, but lacked IFNγ-associated signature (**Figure 2A**).

**Figure 2.**
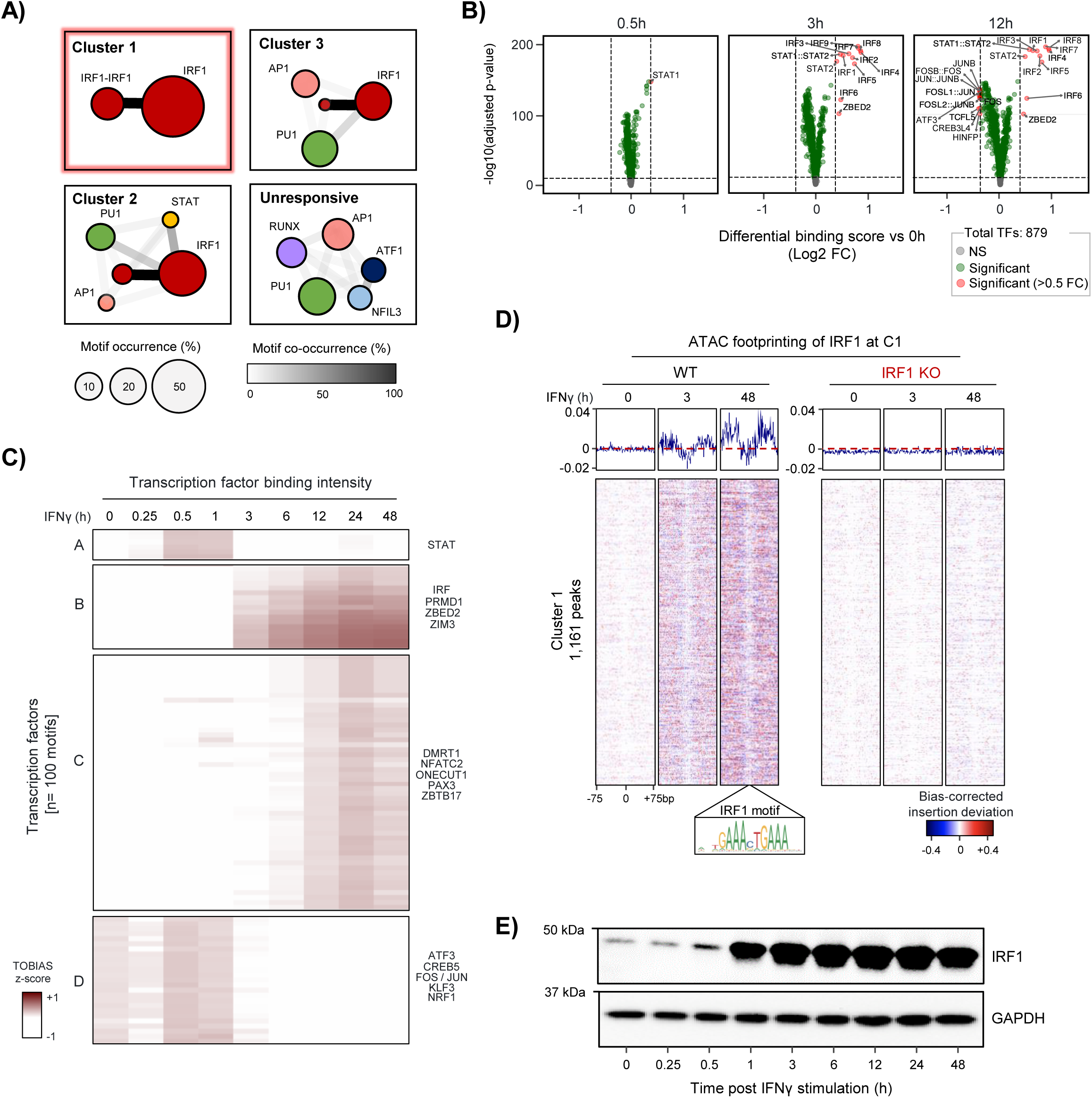
ATAC footprinting and motif network analysis reveal TF activity underpinning IFN-γ response in macrophages. **(A)** Network diagrams of transcription factor motif frequency (node size) and co-occurrence (edge thickness) within ±100 bp of ATAC-seq peak centre for clusters C1-C3 and unresponsive sites. “IRF1–IRF1” denotes sites with ≥2 IRF motifs, and node/edge scales reflect motif frequency and co-occurrence. **(B)** Volcano plots of TOBIAS differential binding scores for 879 mammalian TFs in WT BMDMs comparing 0.5, 3 and 48 h post-IFNγ versus non-treated (0 h); significant TFs are highlighted [Bonferroni-corrected FDR < 0.05; log2 FC > |0.5|]. **(C)** Heatmap of centered TOBIAS TF footprinting intensity in WT BMDMs, grouped into four clusters by k-means clustering. **(D)** Density plots (top) and motif-centered TOBIAS footprint heatmaps (bottom) in WT and IRF1 KO BMDMs showing aggregated IRF1-centered footprinting signal at Cluster 1 sites (rows = individual sites; columns = base position around motif). **(E)** Representative Western blots in WT BMDMs showing IRF1 protein and GAPDH control across time points (0–48 h) post-IFNγ stimulation.

We next conducted differential DNA footprinting on the 38,564 IFNγ-responsive regions to identify the dynamically occupied TF motifs (**Figure 2B**). STAT motifs were the earliest to show differential occupancy (0.5 h), followed by ZBED2, strengthening of STAT, and robust occupancy at IRF motifs by 3 h. At 12 h, FOS/JUN (AP-1) motifs showed disengagement. Clustering the top 100 motifs by TOBIAS binding scores across the time course (0 to 48 h) uncovered four temporal groups: (A) early binding (STAT1), (B) intermediate (IRF, PRDM1, ZBED2, ZIM3), (C) late (DMRT1, NFATC2, ONECUT1, PAX3, ZBTB17), and (D) progressively diminished (ATF3, CREB, FOS, KLF3, NRF1) (**Figure 2C**). These binding dynamics closely mirrored known IFNγ signaling cascades (**Supplementary Figure 2B**).

To directly assess IRF1 motif occupancy at C1 sites, we performed IRF1-centered footprinting at 0, 3, and 48 hours in WT and IRF1 KO macrophages (**Figure 2D**). A characteristic Tn5 protection pattern emerged at IRF1 motifs by 3 h in WT cells and persisted through 48 h, but was completely absent in IRF1 KO cells, suggesting direct and sustained IRF1 binding. Western blot analysis demonstrated IRF1 protein levels were low at baseline, peaked between 3-6 h, and remained elevated through 48 h (**Figure 2E**), paralleling the progressive and persistent motif occupancy (**Figure 2**) and chromatin accessibility observed in WT but not KO macrophages (**Figure 1C**). Collectively, these results suggest that IRF1 is the key regulator orchestrating IFNγ-induced transcriptional responses, with the most enriched and dynamically occupied motif across responsive chromatin.

### Pioneer IRF1 binds closed to establish latent enhancers during macrophage activation

Our motif and footprinting analyses identified IRF1 as the main motif in Cluster 1 (C1), prompting us to directly assess IRF1 binding by ChIP-seq. Mapping the IRF1 ChIP–seq signal onto our ATAC-seq heatmap revealed that IRF1 occupied multiple clusters (C1-3, C8, and unresponsive sites), with the strongest most sustained at C1–C3 following IFNγ stimulation (**Figure 3A, Supplementary Figure 3A**). As control, no signal was detected in IRF1 KO macrophages (**Figure 3A**). At C1 sites, IRF1 binding was detectable by 0.5 h, preceding detectable chromatin accessibility (ATAC–seq) at 3 h, indicating that IRF1 can engage closed chromatin (**Figure 3A, Supplementary Figure 3B**).

**Figure 3.**
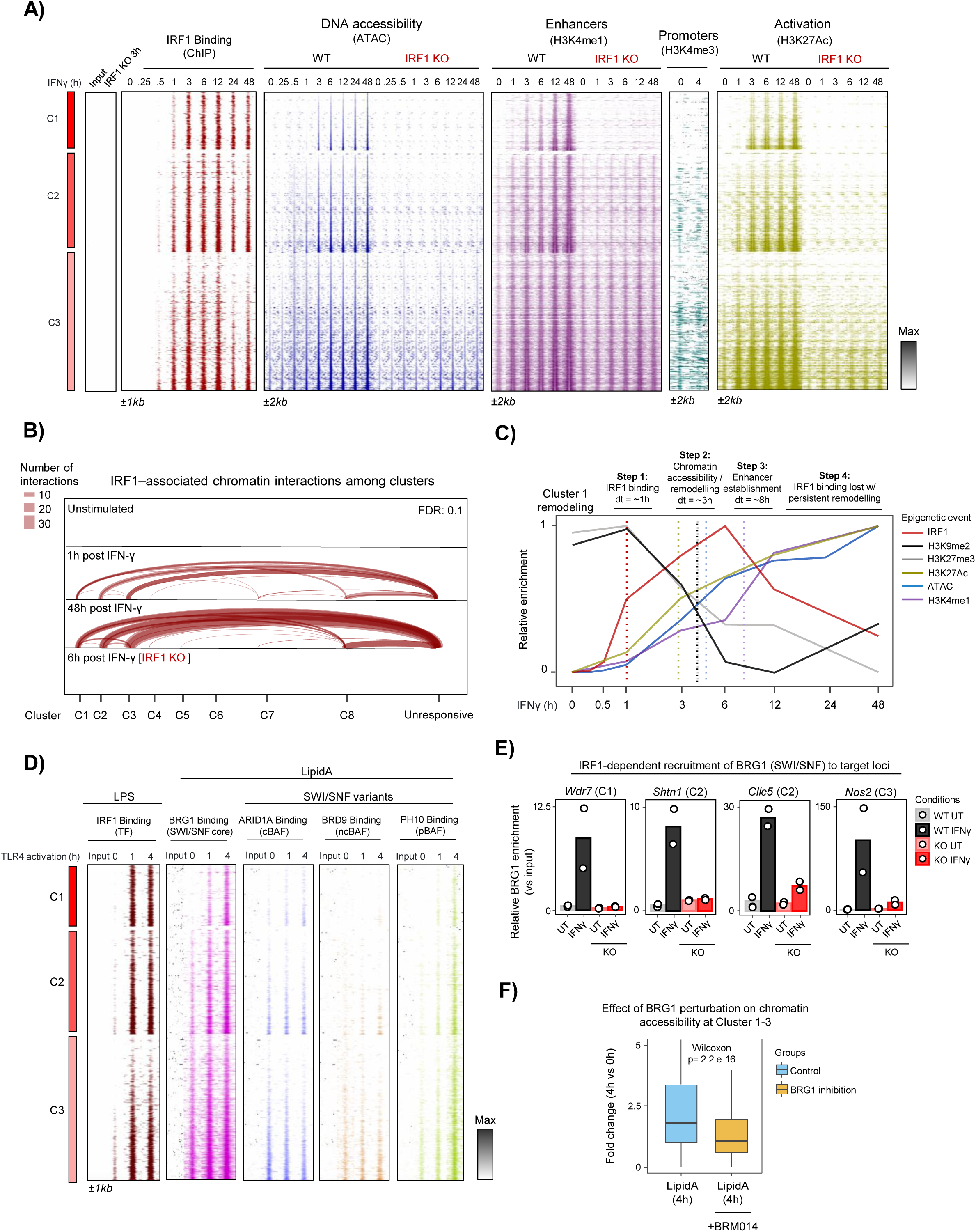
Pioneer IRF1 binds heterochromatin, establishing latent enhancers and recruiting SWI/SNF in response to pro-inflammatory stimuli. **(A)** Heatmaps of IRF1 occupancy (ChIP-seq), ATAC-seq accessibility and H3K4me1, H3K4me3 and H3K27ac signals across in response to IFNγ for sites in Clusters 1–3 in WT and IRF1 KO BMDMs. **(B)** Hi-ChIP arc plots showing loop contacts (arc width represents number of contacts) between IRF1-bound enhancers and promoters at C8 and unresponsive sites in WT and IRF1 KO BMDMs. [FitHiChIP thresholds FDR < 0.1; loop FC > 6, CPM > 6] **(C)** Graph of the temporal changes for ChIP-seq and ATAC-seq signals at Cluster 1. Half-time (t½) to reach 50% of each signal’s maximum was calculated by normalizing each trajectory to its maximum and extracting the pseudo-time at half-max. **(D)** Heatmap of ChIP-seq for IRF1, BRG1, ARID1A, BRD9 and PHF10 across Clusters 1–3 at 0, 1, and 4 h post TLR4 activation. (**E**) BRG1 ChIP–qPCR enrichment (fold over input) at four enhancers (*Wdr7* (C1), *Shtn1* (C2), *Clic5* (C2)*, Nos2* (C3)) in WT and IRF1 KO BMDMs, untreated and 4 h post-IFNγ. **(F)** Boxplots of normalized ATAC-seq counts in Clusters 1–3 in WT BMDMs TLR4 activated with LipidA, with or without and BRG1 inhibition (BRM014).

Next, we defined chromatin states across clusters using ChIP-seq for histone post-translational modifications. H3K4me1 and H3K27ac, marking respectively enhancers and transcriptionally active chromatin, progressively increased at C1–C3 in response to IFNγ (**Figure 3A, Supplementary Figure 3A**). Consistent with *de novo* enhancer formation, C1 sites lacked baseline chromatin accessibility and enhancer-associated marks but acquired ATAC signal concomitantly with H3K4me1 and H3K27ac after stimulation (**Figure 3A, Supplementary Figure 3B**). By contrast, the promoter-associated mark H3K4me3 was absent from C1 sites, and instead enriched at C8, unresponsive regions, and some of C2 and C3 sites (**Supplementary Figure 3A**). The acquisition of ATAC-seq accessibility, H3K4me1, and H3K27ac at C1-3 sites was dependent on IRF1, as was the removal of the repressive mark H3K27me3. Importantly, C1 sites were further distinguished by their localization in facultative heterochromatin, defined by lack of baseline ATAC-seq signal and enrichment of H3K9me2, which was progressively lost upon IFNγ stimulation in a completely IRF1-dependent manner (**Supplementary Figure 3A-B**). Together, these features define C1 as latent enhancers established de novo during macrophage activation.

To determine whether IRF1-dependent enhancer formation translates into functional enhancer–promoter communication, we next examined three-dimensional chromatin interactions using IRF1 Hi-ChIP. Hi-ChIP uncovered extensive looping between IRF1-dependent clusters C1–C3 and unresponsive sites, with comparatively few interactions among themselves or with C8 regions (**Figure 3B**). Notably, unresponsive regions were enriched for H3K4me3 and H3K27ac, indicating that they largely correspond to active gene promoters (**Supplementary Figure 3A**). These findings suggest that IRF1-dependent enhancers in C1–C3 engage pre-existing promoters to drive gene activation. Thus, IRF1-driven latent enhancers are functionally integrated into the macrophage regulatory architecture through long-range chromatin interactions.

To resolve the temporal order of chromatin remodeling, we integrated chromatin features by profiling signal dynamics over the 0 to 48 h time course. C1 latent enhancers underwent a stepwise remodeling process: rapid IRF1 binding (half-time to maximum signal of ∼1 hour), followed by synchronized gains of accessibility and H3K27ac and loss of H3K9me2/H3K27me3 (∼3 h), with further, then accumulation of H3K4me1 (∼8 h), and sustained enhancer maintenance despite declining of IRF1 binding after 12 h (**Figure 3C**). Compared with C2 and C3, C1 displayed slower IRF1 binding and chromatin remodeling rates (**Supplementary Figure 3C–D**). C1 sites required the longest time for full remodeling, followed by C2 and then C3 that remodeled the fastest (**Supplementary Figure 3E**), consistent with their respective baseline chromatin accessibility levels (**Figure 1E**). Together, these data establish IRF1 as a pioneer TF that binds closed chromatin and drives the de novo formation of latent enhancers during IFNγ–induced macrophage activation.

### IRF1 recruits BRG1–SWI/SNF complexes to remodel latent enhancers

Given the extensive chromatin remodeling observed at IRF1-dependent enhancers, we next asked whether IRF1 recruits ATP-dependent chromatin remodelers. Analysis of published datasets (Liao et al., 2024) revealed robust co-recruitment of SWI/SNF components at C1-3 enhancers in BMDMs following Lipid A stimulation, the lipid anchor of lipopolysaccharide (LPS) and agonist of TLR4 (**Figure 3D**). Accordingly, IRF1 ChIP-seq after LPS stimulation (Mancino et al., 2015) showed strong IRF1 recruitment to the C1–3 enhancers, coincident with rapid loading of SWI/SNF subunits (**Figure 3D**): BRG1 binding was detectable within 1 h of stimulation, together with ARID1A (cBAF) and PHF10 (pBAF).

To determine whether SWI/SNF loading depends on IRF1, we performed BRG1 ChIP-qPCR in WT and *Irf1^-/-^* BMDMs treated or not with IFNγ for 4 h (Figure 3E). IFNγ markedly increased BRG1 enrichment at the four tested sites *Wdr7* (C1), *Shtn1* (C2), *Clic5* (C2), *and Nos2* (C3) enhancers in WT cells, whereas recruitment was abolished in IRF1 KO cells, demonstrating IRF1 is required for SWI/SNF targeting (**Figure 3E**). Functionally, pharmacological inhibition of BRG1 with BRM014 significantly blunted Lipid A-induced chromatin accessibility at IRF1-dependent sites after 4 h (**Figure 3F**). These findings indicate that BRG1-containing SWI/SNF complexes are required to achieve full IRF1-driven chromatin opening. Ultimately, the interplay between IRF1 and SWI/SNF complexes enables stable, accessible regulatory elements, providing a mechanistic framework for how IRF1 modulates macrophage activation through chromatin reprogramming.

### High-density motif arrays underpin IRF1 pioneering activity

We profiled 55,416 genome-wide IRF1 binding sites in BMDMs from 0 to 48 h post-IFNγ stimulation to understand IRF1 occupancy dynamics. Sites were ranked by mean IRF1 ChIP-seq signal and stratified into four occupancy classes: weak, moderate, strong, and very strong (**Supplementary Figure 4A**). All classes showed IFNγ-induced IRF1 recruitment (Figure 4A), but the very strong sites (top 10%) displayed >10-fold higher IRF1 signal relative to the other classes (**Figure 4B**). Notably, these very strong sites were characterized by low baseline accessibility and underwent progressive chromatin opening following IFNγ, whereas weak sites remained largely inaccessible. Moderate and strong sites were mostly pre-accessible before IFNγ but some sites still required IRF1 to gain full accessibility (**Supplementary Figure 4B**). Mapping these IRF1 classes to our previously defined ATAC clusters revealed that clusters C1–C3 predominantly intersected with the very strong IRF1 sites, whereas the other IRF1 binding classes tended to fall into clusters C4–C8 or unresponsive regions (**Figure 4C, Supplementary Figure 3A**). Strikingly, ∼50% of IRF1-binding events occurred outside ATAC clusters (**Supplementary Figure 4C**), consistent with low-level sampling of nucleosomal DNA, a hallmark of other pioneer factors (Lerner et al., 2023).

**Figure 4.**
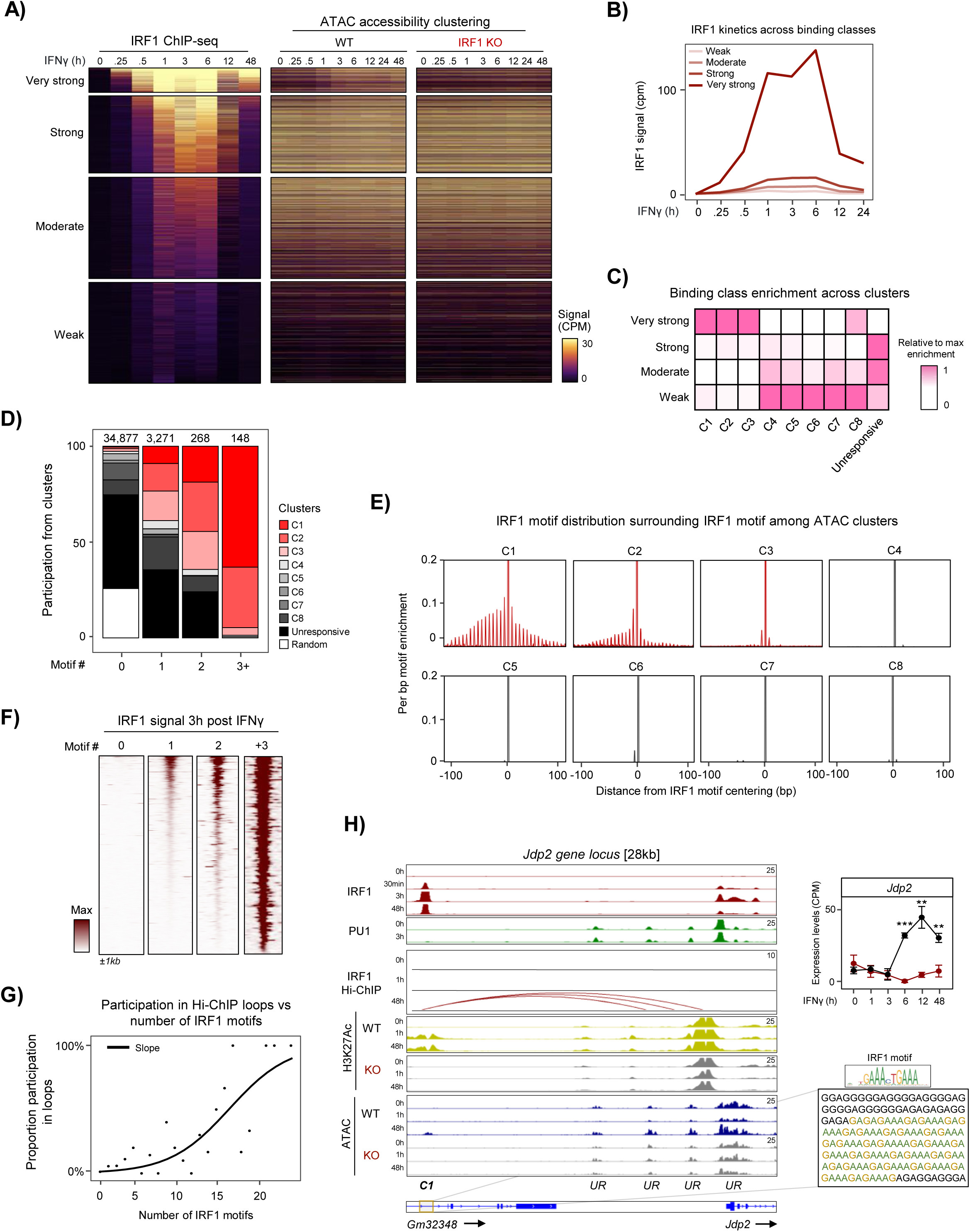
TF concentration and high-density motif arrays underpin IRF1 pioneer activity. **(A)** Heatmaps of IRF1 ChIP–seq and normalized ATAC–seq at sites grouped by IRF1 signal strength (very strong to weak) in response to IFNγ. **(B)** Line plots of average IRF1 ChIP–seq signal in WT BMDMs for each binding-strength category. **(C)** Heatmap of relative enrichment of IRF1 binding classes across ATAC clusters (enrichment is relative to the maximum site overlap). **(D)** Stacked bar plots showing proportions of sites with 0, 1, 2 or ≥3 IRF1 motifs per ATAC cluster. **(E)** Aggregate plots of IRF1 motif frequency across ±100 bp around IRF1 peaks for each ATAC cluster. **(F)** Heatmap of IRF1 ChIP–seq signal at 3 h post–IFNγ for sites stratified by IRF1 motif count, as determined in D). **(G)** Scatter plot of fraction of sites forming IRF1 Hi-ChIP loops versus motif count, with a fitted trend line shown. [FitHiChIP thresholds FDR < 0.1; loop FC > 6, CPM > 6] **(H)** Genome browser tracks at the *Jdp2* locus showing IRF1, PU.1 and H3K27ac ChIP–seq, Hi-ChIP interactions and ATAC–seq in WT and IRF1 KO BMDMs. The cluster to with each ATAC-seq peak belong is indicated [C1 = cluster 1; UR = Unresponsive]. Insets display the array of IRF1 motifs at the C1 site and a SLAM-seq *Jdp2* expression plot across the IFNγ time course.

We next asked whether IRF1 occupancy level was dictated by motif number and architecture. We reasoned that the enhanced pioneering at very strong sites might be driven by the presence of multiple IRF1 motifs. Indeed, sites harboring higher numbers of IRF1 motifs were strongly enriched in C1–C3, with the highest enrichment in C1 (**Figures 4D**). Interestingly, when centering on one IRF1 motif and scanning for additional motifs, we found that in C1 sites, and to a lesser extent in C2 sites, supplemental IRF1 motifs were regularly interspaced, suggesting an ordered motif architecture (**Figure 4E**). De novo motif analyses further confirmed the presence of high-density IRF1 motif arrays in C1 (**Supplementary Figure 4D**). IRF1 ChIP-seq signal scaled with motif number, supporting a direct relationship between motif density and higher IRF1 occupancy (**Figure 4F**). In IRF1 KO macrophages, multi-motif sites completely failed to acquire ATAC accessibility or did not undergo removal of repressive marks (H3K27me3, H3K9me2), underscoring IRF1’s requirement for chromatin remodeling (**Supplementary Figure 4E**). Activation of these sites, as marked by deposition of H3K27ac, was also completely abrogated, including at sites containing only one or two IRF1 motifs, indicating a strict dependency on IRF1 recruitment for enhancer activation. Hi-ChIP analysis further revealed that enhancer-promoter looping frequency increased with IRF1 motif number, reaching a plateau approximately at 15 motifs (**Figure 4G; Supplementary Figure 4F**). As an illustrative example, the *Jdp2* locus contains a C1 enhancer with a dense IRF1 motif array that displayed strong IFNγ-induced IRF1 binding, gain of accessibility and H3K27ac deposition, and Hi-ChIP interactions in WT cells, but remained closed and transcriptionally silent in KO macrophages (**Figure 4H**).

Taken together, these data demonstrate that IRF1 pioneering activity is quantitatively encoded by motif density and TF occupancy level. Dense IRF1 motif arrays promote higher IRF1 occupancy, enable chromatin remodeling at otherwise inaccessible loci, and promote productive long-range enhancer-promoter interactions.

### IRF1 drives enhancer-linked transcriptional rewiring in activated macrophages

We next evaluated how IRF1-driven enhancers impacts transcription by performing a SLAM-seq time course (0–48 h post-IFNγ) to quantify the total and nascent transcriptome in WT and IRF1 KO BMDMs (**Figure 1A**). Principal component analysis (PCA) revealed a striking separation between the trajectories of transcriptional changes to IFNγ in WT and IRF1 KO macrophages. Nascent reads captured most of the IFNγ-induced transcriptional variance with only 8% of total reads (**Supplementary Figure 5A**). To link enhancer dynamics with transcriptional output, we quantified nascent transcription of genes nearest each ATAC-defined enhancer cluster. Gene expression trajectories closely mirrored chromatin accessibility kinetics (**Figures 5A, 1C**), with clusters C1–C3 showing robust IFNγ-induced transcription in WT cells that was markedly blunted in IRF1 KO macrophages. As for the ATAC signal, C5 proximal genes showed increased response in IRF1 KO cells, likely through secondary mechanism since IRF1 is not recruited to these sites. The other clusters displayed IRF1-independent changes matching their accessibility trajectories; for example, C6 exhibiting transient and C8 late transcriptional activation. Together, these results establish a direct coupling of IRF1-dependent activation and transcriptional modulation at nearby loci.

**Figure 5.**
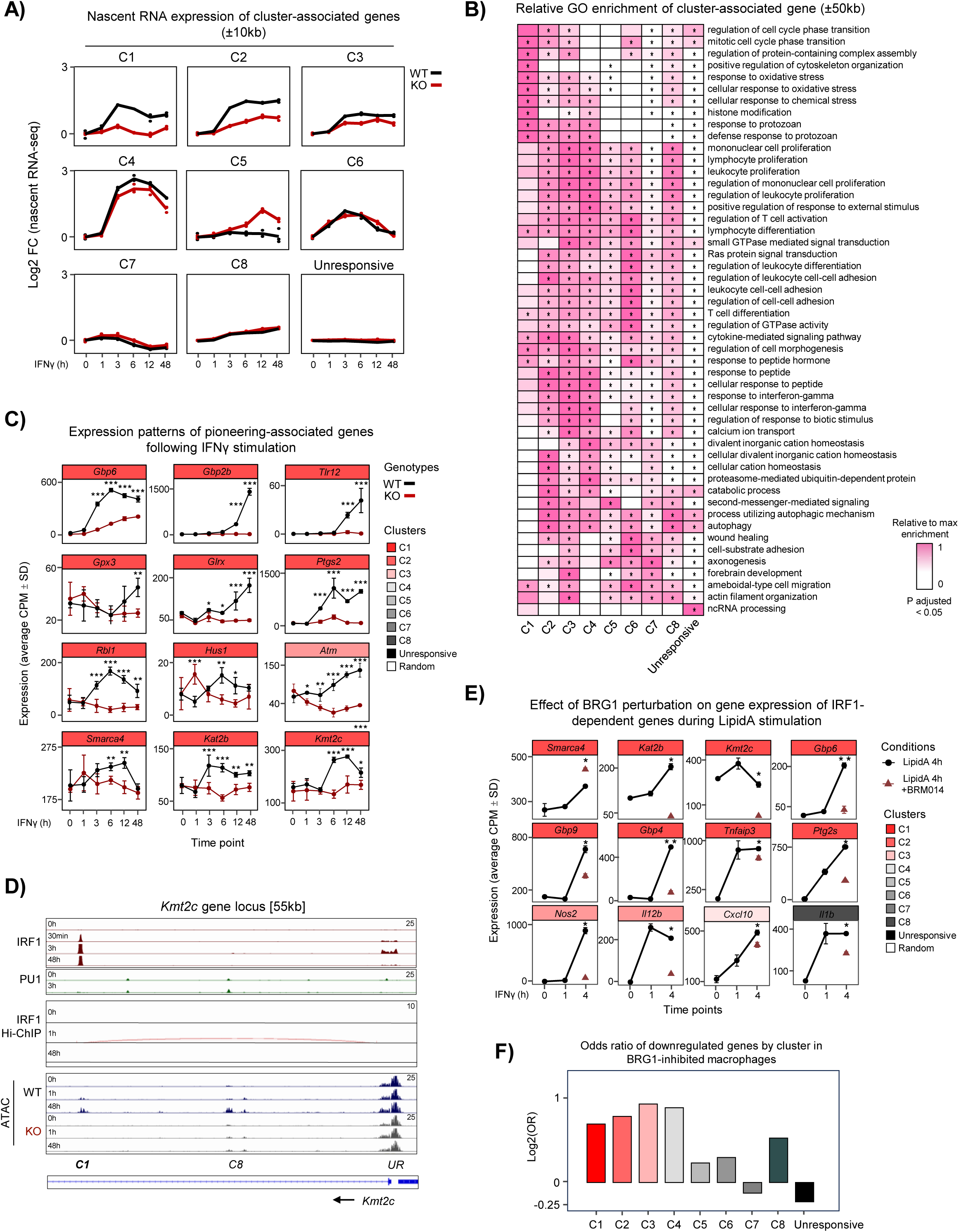
IRF1 pioneers non-canonical gene programs through the IRF1-BRG1 axis. **(A)** Line plots of nascent RNA-seq log2 fold-change (FC) for genes within ±10 kb of ATAC cluster regions in WT and IRF1 KO BMDMs in response to IFNγ stimulation. **(B)** Heatmap of GO biological process enrichment for genes within ±50 kb of ATAC peaks. Categories with clusterProfiler FDR < 0.05 for at least one cluster are shown. **(C)** Line plots of nascent RNA counts per million (CPM; mean ± SD) for selected genes across IFNγ time points; WT vs IRF1 KO comparison by two-way ANOVA and pairwise post-hoc testing at each time point. **(D)** Genome browser tracks at the *Kmt2c* locus showing IRF1 and PU.1 ChIP-seq, Hi-ChIP arcs and ATAC-seq signal for WT and IRF1 KO BMDMs. [UR = Unresponsive] **(E)** Line plots of RNA-seq CPM (mean ± SD) for selected genes at 0, 1 and 4 h post-LipidA treatment in WT BMDMs, with BRM014 treatment at the 4 h time point. [Student T-test; n = 3] **(F)** Bar plot of log2 odds ratio of downregulated genes (FC < 0.5 and FDR < 0.05) after BRM014 treatment (4 h Lipid A) across clusters. * < 0.05, ** < 0.01, *** < 0.001

IRF1 appears to build enhancers *de novo*—an energetically costly and time-consuming investment for the cell. To understand the purpose of these newly forged elements, we next asked which biological programs they control. We therefore performed comparative GO analysis across all enhancer clusters by assigning each ATAC-seq peak to its nearest gene. As depicted on the relative enrichment heatmap (**Figure 5B**), genes associated to IFNγ-induced clusters are enriched in functions associated to innate immune responses and activation of adaptive immunity, as expected from macrophage activation. Interestingly, genes associated with the latent enhancer C1 cluster were selectively enriched for pathways linked to defense response to protozoan, histone modifications, oxidative stress response and regulation of cell cycle transition. Consistently, the expression of many genes within these pathways was severely impaired in IRF1 KO BMDMs. In line with IRF1’s immunological roles, canonical immune effectors such as *Gbp6*, *Gbp2b* and *Tlr12* were robustly induced in WT but reduced in IRF1 KO cells (**Figure 5C**). Unexpectedly, IRF1-dependent latent enhancers also controlled multiple chromatin remodelers and epigenetic regulators, including *Smarca4* (encoding BRG1), *Kat2b* (encoding the histone acetyltransferase PCAF), and *Kmt2c* (the H3K4me1 methyltransferase that establishes enhancer marks), were significantly upregulated in WT macrophages but not in IRF1 KO cells (**Figure 5D**). An unbiased analysis of genes included in the GO category “GOBP Chromatin Remodeling” further highlighted a group of IRF1-dependent modifiers such as *Hdac9*, *Kat2a* (GCN5), *Kat5* (TIP60), *Kmt2b*, Smarca5 (ISWI), and *Chd1* (**Supplementary Figure 5C**). For example, at the *Kmt2c* locus (**Figure 5D**), IRF1 bound a distal regulatory site lacking PU.1 occupancy that became accessible exclusively in WT cells. Hi-ChIP data confirmed IRF1-dependent looping between this newly established enhancer and the *Kmt2c* promoter as early as 1 h post-stimulation. These findings suggest that IRF1 initiates a feed-forward transcriptional program that reinforces and amplifies epigenetic reprogramming during macrophage activation.

Beyond immune effector genes, IRF1 controlled genes linked with cell cycle regulation and DNA damage, as well as oxidative stress management. For example, oxidative stress–related genes (*Gpx3*, *Glrx*, *Ptgs2*) and DNA damage checkpoint genes (*Rbl1*, *Hus1*, *Atm*) exhibited robust IRF1-dependent induction (**Figure 5C**). At the *Rbl1* locus, IRF1 bound a distal regulatory site devoid of PU.1, accompanied by IRF1-dependent chromatin remodeling and long-range promoter contacts at late time points (**Supplementary Figure 5D**). Gene set enrichment analysis (GSEA) further indicated coordinated attenuation of p53-related stress response pathways, such as “Reactome_Stabilization_of_P53”, in IRF1 KO macrophages, consistent with impaired stress adaptation during sustained inflammatory signaling (**Supplementary Figure 5E**).

Given the IRF1-dependent recruitment of BRG1 to C1 enhancers and the impaired chromatin accessibility observed upon SWI/SNF inhibition (**Figures 3E-F**), we postulated that SWI/SNF activity is required for IRF1-driven transcriptional programs. Analysis of published datasets BRG1-inhibited BMDMs during Lipid A stimulation (Liao et al., 2024), revealed selective loss of activation of key pioneer targets, including *Smarca4*, *Kmt2c*, and *Gbp6* (**Figure 5E**). Moreover, odds ratio analysis confirmed a disproportionate blockade of activation among IRF1-dependent clusters (**Figure 5F**). Lastly, we confirmed these inhibition patterns during IFNγ stimulation. Indeed, BRG1 inhibition blocked the IRF1-dependent IFNγ response of the tested genes (**Supplementary Figure 5F**).

Together, these results demonstrate that IRF1-dependent enhancer formation drives a broad transcriptional reprogramming program in activated macrophages, coupling immune effector functions, chromatin remodeling, and cellular stress responses. The dependence of these programs on BRG1-containing SWI/SNF complexes underscores a mechanistic axis through which IRF1 pioneers’ non-canonical transcriptional states during inflammation.

### IRF1 orchestrates metabolic reprogramming for macrophage effector function

IFNγ is known to reprogram macrophage metabolism by promoting aerobic glycolysis, pentose phosphate pathway (PPP), altering the tricarboxylic acid (TCA) cycle, and suppressing oxidative phosphorylation (OXPHOS) (O’Neill & Pearce, 2016; Wang et al., 2018). Building on this framework, we tested whether IRF1 directly mediates the metabolic rewiring that follows IFNγ stimulation.

Genome-wide IRF1 ChIP-seq revealed extensive IRF1 binding at genes encoding metabolic enzymes and regulators of glycolysis, the PPP, and the TCA cycle (**Figure 6A**). At the *Hk1* locus—encoding hexokinase 1 which mediates the initial step of glycolysis by catalyzing phosphorylation of D-glucose to D-glucose 6-phosphate—IRF1 occupied and opened an intragenic enhancer (C2 site), with Hi-ChIP chromatin contact maps indicating IRF1-dependent enhancer–promoter coupling (**Figure 6B**) maintenance of increased *Hk1* expression in response to IFNγ (**Figure 6C**). Consistently, transcriptomic profiling revealed widespread IRF1-dependent dysregulation of metabolic genes following IFNγ, affecting glucose transporters, glycolytic enzymes, PPP components and TCA cycle factors (**Figure 6C; Supplementary Figure 6A**). To evaluate the functional consequences, we next measured oxygen consumption rates (OCR) over 24 h. At steady state, WT and IRF1 KO BMDMs showed comparable OCR (**Figure 6D**). Following IFNγ stimulation, WT macrophages exhibited a marked reduction in oxidative consumption beginning ∼3 h, whereas IRF1 KO cells failed to downregulate OXPHOS (**Figure 6D**), indicating a defect in metabolic switching. IRF1 KO macrophages also displayed higher OCR than WT cell following LPS treatment, but not in response to β-glucan (BDG), although both stimuli induced only modest changes in OCR overall (**Supplementary Figure 6B**).

**Figure 6.**
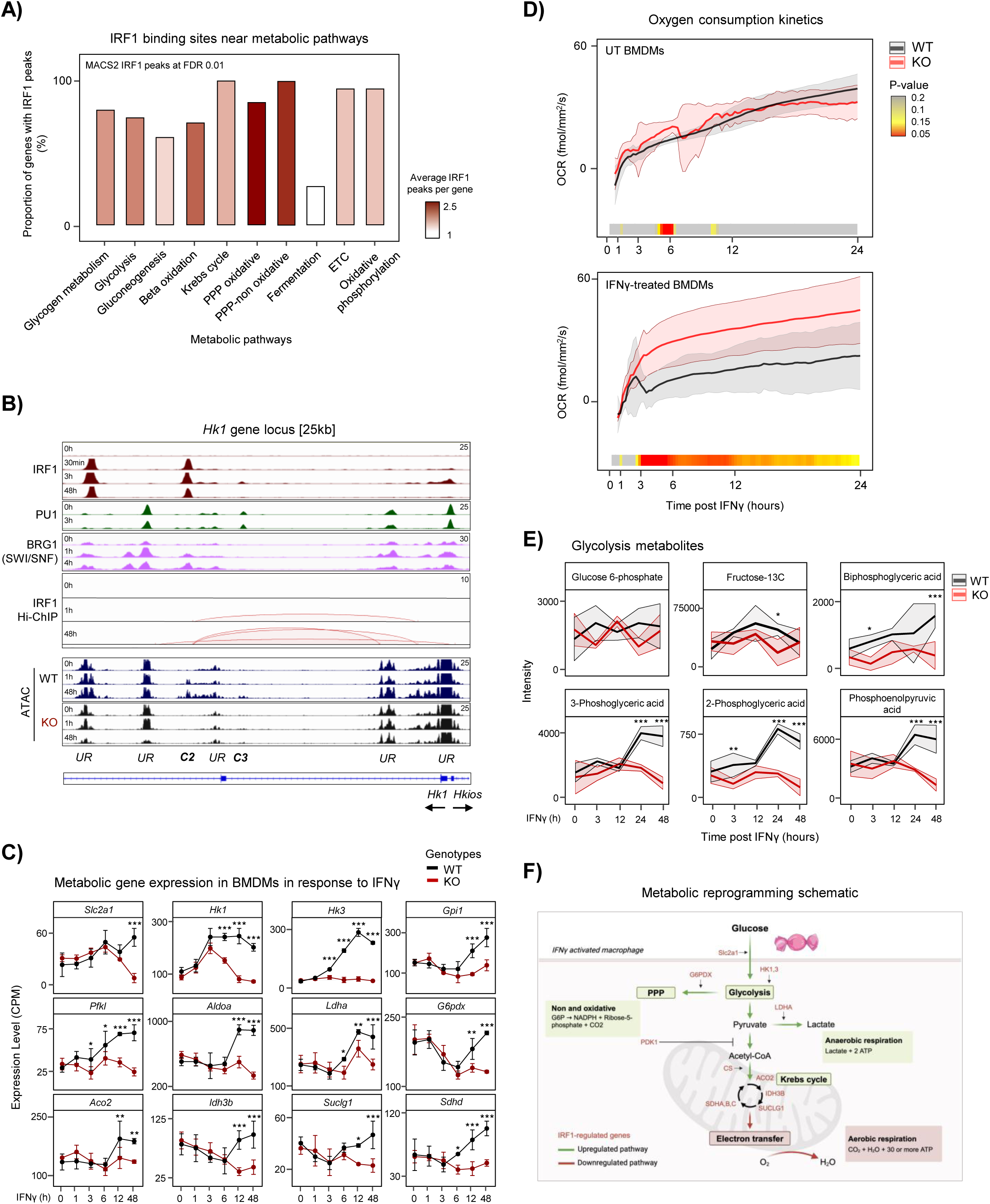
IRF1 governs metabolic shift into glycolysis to sustain macrophage activation. **(A)** Bar plot of the proportion of genes in selected metabolic pathways that harbor IRF1 ChIP-seq peaks; red intensity denotes average number of peaks per gene in each pathway. **(B)** Genome browser tracks of the *Hk1* locus showing normalized IRF1 and PU.1 ChIP-seq, Hi-ChIP interactions and ATAC-seq; an inset shows the annotated intragenic enhancer and promoter contact. [UR = Unresponsive] **(C)** Line plots of nascent RNA CPM (mean ± SD) for selected genes in glycolysis, PPP and TCA pathways WT and IRF1 KO BMDMs; two-way ANOVA and post-hoc testing; * < 0.05, ** < 0.01, *** < 0.001. **(D)** Oxygen consumption rates (OCR; fmol mm⁻² s⁻¹) for untreated and IFNγ–stimulated WT and IRF1 KO BMDMs [n = 4/group]; adjacent heatmap shows Student t-test p-values for each time point measured. (**E**) Ribbon plots of relative glycolysis metabolite intensity (mean ± SD) detected by GC-MS in response to IFNγ in WT and IRF1 KO BMDMs [n = 3/group]. **(F)** Diagram of glycolysis, pentose phosphate pathway (PPP) and Krebs cycle highlighting genes and pathway components significantly dysregulated in IRF1 KO BMDMs for at least 1 timepoint.

Targeted gas chromatography (GC) and liquid chromatography (LC) coupled with mass spectrometry (MS) further revealed increased abundance in several glycolytic intermediates in WT but not in IRF1 KO macrophages (**Figure 6E**). Notably, the metabolites elevated in WT cells, including 3-phosphoglycerate, 2-phosphoglycerate, 2,3-bisphosphoglycerate, and phosphoenolpyruvate, map to the later, energy-generating phase of glycolysis, consistent with previous reports that IFNγ increases glycolytic flux (Wang et al., 2018), which we now describe being downstream and dependent on IRF1. The accumulation of these late glycolytic intermediates is also consistent with increased diversion of upstream carbon into the PPP, supporting NADPH production and redox balance during macrophage activation. Together, these data position IRF1 at the key mechanistic bridge between inflammatory signalling and glycolytic reprogramming in IFNγ-activated macrophages (**Figure 6F**).

A global differential abundance analysis comparing wild type at 48 hours to 0 hours identified itaconic acid, succinic acid and mesaconic acid among the most strongly upregulated metabolites upon IFNγ stimulation (**Figure 7A, left**). Direct comparison of WT and IRF1 KO macrophages at 24 h confirmed that accumulation of itaconic acid and increases in several Krebs cycle intermediates were IRF1 dependent (**Figure 7A, right**). Focused analysis of the pentose phosphate pathway showed lower levels of sedoheptulose-7-P, xylulose and erythrose in IRF1 KO cells (p = 3.29e-2, p = 1.77e-3 and p = 4.71e-6, respectively) (**Figure 7B**). Moreover, IRF1 binding was detected at the *G6pdx* promoter by 3 h post-IFNγ stimulation (**Supplementary Figure 7A**), as well as IRF1-dependent gene expression at later time points (**Figure 6C**). Indicative of the decreased PPP output in IRF1 KO, ADP-ribose, a PPP-associated metabolite and a proxy for NAD⁺ consumption, was significantly lower in KO at 48 h (p = 0.0075; **Supplementary Figure 7B**). Consistently, redox profiling revealed an increased GSH:GSSG ratio in WT relative to IRF1 KO, with divergence emerging as early as 3 h and reaching significance at 24 h (p = 0.0037) (**Figure 7C; Supplementary Figure 7C**).

**Figure 7.**
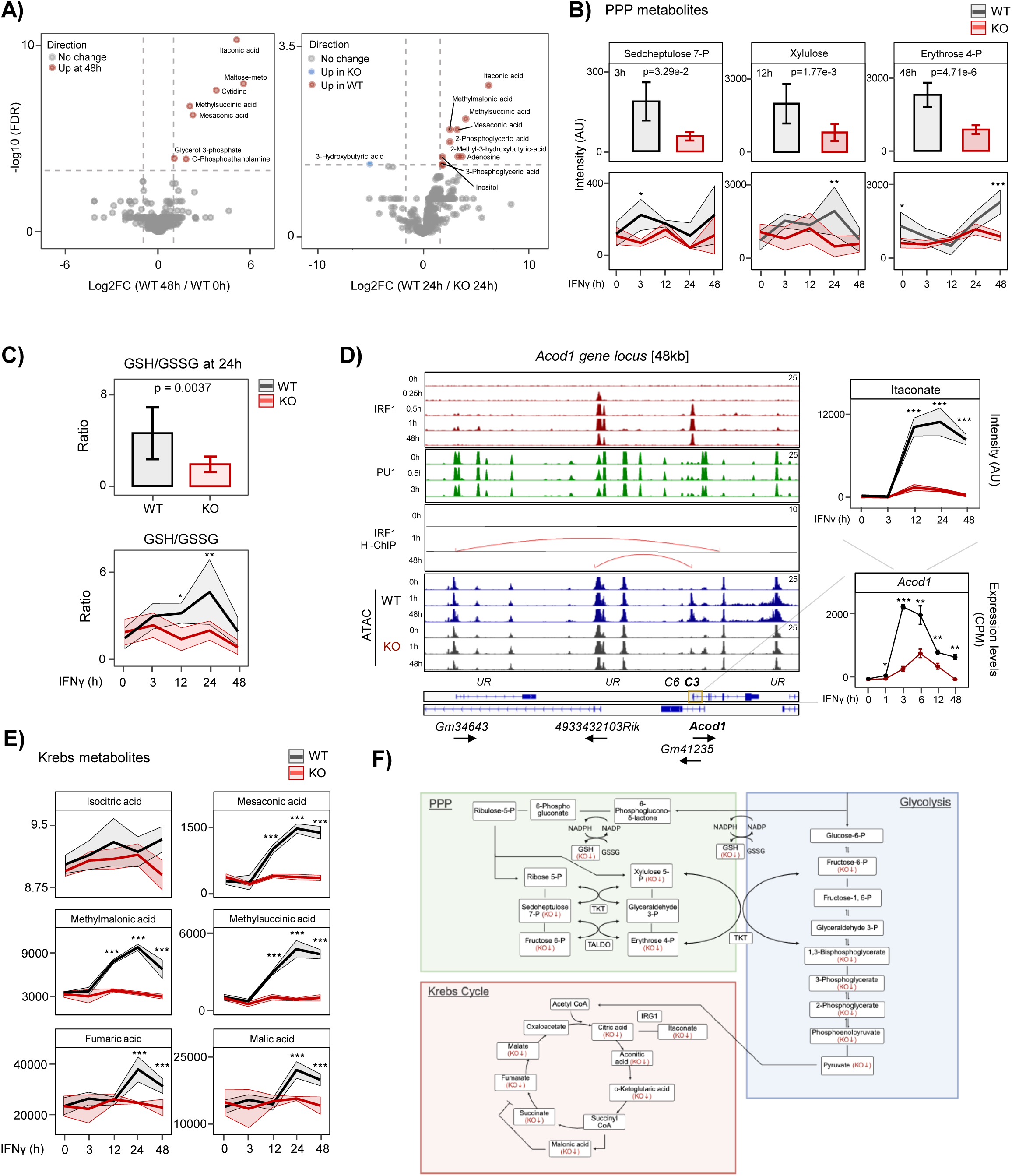
IRF1 redirects glycolytic carbon into the pentose phosphate and TCA cycles to sustain redox and immunoregulatory metabolism. **(A)** Volcano plots from differential metabolite abundance analysis for GC-MS data (n=3/group), comparing 48 h versus 0 h in WT cells (left) and WT versus IRF1 KO at 48 h (right). **(B)** Top: bar plots of normalized GC-MS intensity for sedoheptulose 7-P at 3 h post IFNγ, xylulose at 12 h, and erythrose 4-P at 48 h. Bottom: ribbon plots of normalized MS signal over time with mean ± SD. **(C)** Top: normalized LC-MS GSH intensity at 24 h post-IFNγ stimulation. Bottom: ribbon plots of GSH/GSSG ratios over time (mean ± SD) calculated from normalized LC-MS intensities [n=3/group]. **(D)** Genome browser tracks at the *Acod1* locus showing normalized IRF1 and PU.1 ChIP-seq, Hi-ChIP interactions and ATAC-seq [UR = Unresponsive]. Adjacent panels show *Acod1* nascent RNA expression and itaconic acid levels. **(E)** Ribbon plots of normalized GC-MS signal for TCA metabolites in response to IFNγ in WT and IRF1 KO BMDMs. **(F)** Diagram of glycolysis, PPP and TCA cycle metabolic pathways with dysregulated intermediates denoted in red.

IRF1 also directly regulated remodeling of the TCA cycle. IRF1 bound the *Acod1* promoter within 15 minutes of IFNγ and chromatin looping was observed at these sites (**Figure 7D**); *Acod1* mRNA was very strongly upregulated in WT in comparison with KO cells which had minimal and delayed induction (**Figure 7D**). Mass spectrometry confirmed parallel, IRF1-dependent accumulation of itaconic acid, an important immunometabolite that links antimicrobial defense with redox and inflammatory control. Additional TCA metabolites, including isocitrate, mesaconic acid, methylmalonic acid, methylsuccinic acid, fumarate and malate, were rapidly induced after IFNγ in an IRF1-dependent manner (**Figure 7E**). Consistent with these changes, IRF1 occupancy was detected at the promoter of *Sdhd*, the gene encoding succinate dehydrogenase, after 3 h post-IFNγ followed by an IRF1-dependent transcriptional increase at 6h and onwards (**Supplementary Figure 7D, Figure 6C**), supporting and altered succinate to fumarate conversion and the observed increase in fumarate. A schematic summary highlights the IRF1-dependent increases in glycolytic intermediates, enhanced PPP engagement and coordinated remodeling of TCA cycle metabolites (**Figure 7F**). Together with the IRF1-dependent induction of itaconate and other immunomodulators, these results demonstrate that IRF1 is required for the coordinated epigenetic, transcriptional, and metabolic reprogramming that characterizes IFNγ-activated macrophages.

### Irf1 is required for long-term enhancer reprogramming of defense and metabolic genes in macrophages

Building on our finding that IRF1-dependent chromatin remodeling persists even after diminished IRF1 binding declines (**Figure 3C**), we investigated whether IRF1’s pioneering activity establishes durable chromatin states. We then performed a wash-and-rest experiment to assess the persistence of enhancer reprogramming following IFNγ withdrawal (**Figure 8A**). BMDMs were stimulated with IFNγ for 24 h, followed by either a 48 h washout period, or a six-day washout, with or without an additional 1 h re-stimulation to probe recall. ChIP-seq for H3K4me1 (enhancer mark), H3K27ac (activation mark) and H3K9me2 (facultative heterochromatin mark) was then performed on the seventh day after plating, to determine whether IRF1-induced chromatin modifications persist in the absence of continuous cytokine signaling.

**Figure 8.**
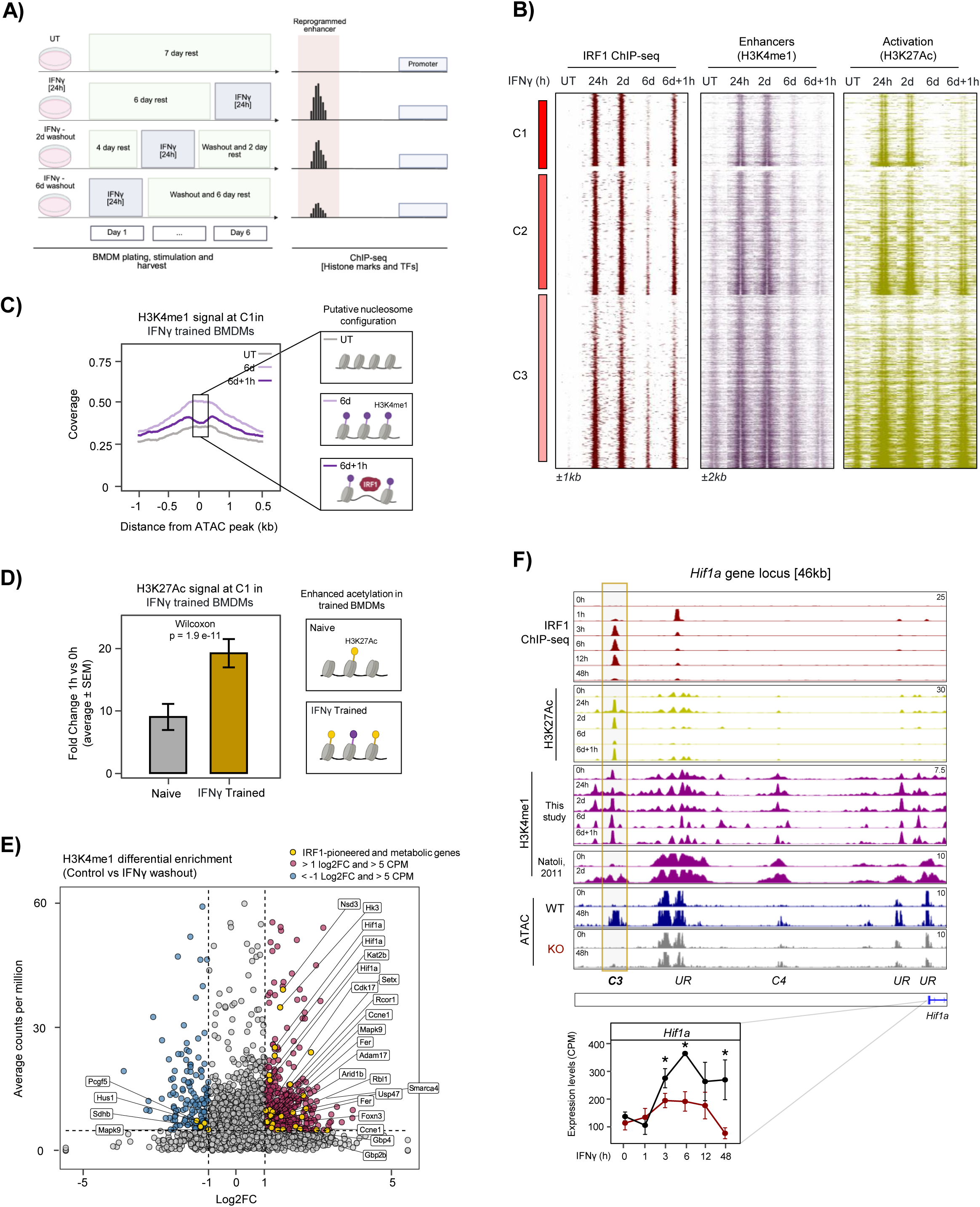
IRF1-primed enhancer memory enables swift re-activation of defense and metabolic loci in macrophages. **(A)** Schematic of experimental timeline for the long-term wash-and-rest assay. Cells are plated for seven days, pulsed with 24 h IFNγ (400 U/mL) at specified times (24 h, 48 h, 6 d) with defined washout intervals and a final 1 h re-stimulation. On day 7, cells are harvested for ChIP-seq (IRF1, H3K4me1, H3K27ac and H3K9me2). **(B)** Heatmaps of normalized ChIP-seq signal for IRF1, H3K4me1, and H3K27ac at Clusters 1–3. **(C)** Aggregate coverage plots of H3K4me1 ±1 kb from ATAC peak centers for UT, 24 h IFNγ, 6 d washout and 6 d + 1 h restimulation; insets show putative nucleosomal configurations. **(D)** Bar plots of fold-change in H3K27ac (mean ± SEM) comparing naïve and IFNγ-trained cells after 1 h restimulation; statistical comparison using Wilcoxon test. **(E)** Volcano plot of H3K4me1 differential enrichment for Cluster 1–3 (control versus IFNγ washout); points = enhancers, color key: red = increased, blue = decreased, yellow = pioneered genes; labeled enhancers meet log₂FC > 1 and CPM > 5. **(F)** *Hif1a* locus showing normalized IRF1, H3K27ac, and H3K4me1 ChIP-seq, and ATAC-seq in WT and IRF1 KO BMDMs [UR = Unresponsive]. Normalized SLAM-seq nascent RNA expression for *Hif1a* is shown; * p < 0.05.

Heatmaps of histone modifications revealed that IRF1-dependent, IFNγ-responsive clusters C1–C3 retained H3K4me1 signal after six days of washout but returned to baseline activity as depicted by H3K27ac signal (**Figure 8B, Supplementary Figure 8B**). Concomitantly, H3K9me2 levels declined across IRF1-bound peaks during the washout (**Supplementary Figure 8A**). Notably, C1 latent enhancers, which lacked pre-existing enhancer marks, acquired H3K4me1 marks in response to IFNγ and maintained it after two and six days of washout (**Figure 8B-C**). Upon the 1 h re-stimulation, these trained enhancers rapidly reacquired H3K27ac (Wilcoxon p = 1.9 × 10⁻¹¹ vs primary response), underscoring their heightened responsiveness (**Figure 8D**). Genome-wide differential analysis between control and washout conditions confirmed persistent H3K4me1 reprogramming at enhancers adjacent to genes involved in metabolism (e.g. *Hk3*, *Hif1a*), chromatin remodeling (*Kat2b*), cell cycle control (*Rbl1, Ccne1*) and host defense (*Gbp2b*, *Gbp4*). We validated these findings at two representative loci. At the guanylate-binding-protein (*Gbp*) locus, where Hi-ChIP showed that IRF1-dependent enhancer–promoter loops are established and remain detectable for at least 48 h, H3K4me1 enrichment at multiple *Gbp* genes persisted after washout (**Supplementary Figure 8C**). Likewise, at the *Hif1a* locus (**Figure 8F**), an IRF1-dependent C3 site retained elevated H3K4me1 six days post-stimulation; consistent with this, nascent *Hif1a* transcription remained IRF1-dependent throughout prolonged IFNγ exposure (0–48 h).

Finally, live-cell oxygen consumption measurement demonstrated that WT BMDMs maintain lower respiratory rate than IRF1-knockout cells during prolonged IFNγ or LPS treatment (**Supplementary Figure 8D**), indicating that IRF1 supports sustained metabolic rewiring away from oxidative phosphorylation under ongoing inflammatory cues.

Together, these data demonstrate that IRF1’s pioneering function established a stable epigenetic “memory” at previously inactive enhancers, characterized by persistent H3K4me1 deposition, loss of repressive H3K9me2, and accelerated re-acetylation upon re-challenge. This durable enhancer priming provides a mechanistic basis for the enhanced defense and metabolic responsiveness of macrophages following IFNγ exposure, highlighting IRF1 as a central determinant of innate immune plasticity.

## DISCUSSION

Our study identifies Interferon Regulatory Factor 1 (IRF1) as an essential regulator of macrophage activation to IFNγ, acting through direct and durable remodeling of the enhancer landscape. Although the lineage-determining pioneer factor PU.1 has long been viewed as the primary architect of macrophage identity (Feng et al., 2008; Minderjahn et al., 2020), we show that a substantial subset of IFNγ-responsive enhancers relies on IRF1, rather than PU.1, for initial accessibility. IRF1 binds rapidly to these regions following stimulation, initiating removal of repressive histone marks (H3K9me2, H3K27me3), chromatin remodeling (ATAC-seq gain), and subsequent deposition of activating marks (H3K4me1, H3K27ac). Notably, while IRF1 binding declines after 12 h, the remodeled enhancers persist long after IFNγ withdrawal, underscoring IRF1’s capacity to induce durable epigenetic reprogramming.

These findings place IRF1 within the framework of latent enhancer activation, in which previously inaccessible, nucleosome-occluded regions become functionally engaged upon inflammatory stimulation (Glass & Natoli, 2016; Ostuni et al., 2013). While earlier work emphasized cooperation between signal-dependent TFs and PU.1 in unmasking latent enhancers after LPS stimulation (Ostuni et al., 2013), our data reveal that IRF1 can also independently pioneer enhancer activation at sites lacking PU.1 occupancy, thereby extending the plasticity of the macrophage epigenome beyond lineage-imposed constraints. Together, these findings clarify prior reports (Karwacz et al., 2017; Song et al., 2021) and establish IRF1 as a bona fide pioneer factor in activated macrophages.

Pioneer transcription factors are distinguished by their ability to engage nucleosomal DNA within compact chromatin and initiate chromatin opening to facilitate subsequent binding of secondary TFs (Frederick et al., 2023; Mayran et al., 2018; Zaret & Carroll, 2011). Our high-resolution temporal analyses revealed that IRF1 binding at latent enhancers (C1 *de novo*, C2 primed, C3 pre-accessible) precedes chromatin accessibility and histone modification changes, mirroring classical pioneer behavior exemplified by FOXA1 and others (Brennan et al., 2023; Mayran et al., 2018; Sinha et al., 2023). Importantly, newly established enhancers require extended remodeling time (up to 24 h), in comparison to primed or pre-accessible enhancers (C2 ∼18 h and C3 ∼12 h, respectively), to fully transition from silent chromatin to an active enhancer state, highlighting the energetic and temporal costs of de novo enhancer formation.

IRF1 pioneering activity is strongly influenced by motif density and transcription factor concentration, with dense IRF1 motif arrays driving higher occupancy, stronger chromatin opening, and enhancer–promoter communication. This mechanism parallels recent findings for OCT4 and SOX2, where adjacent motifs synergize to enhance local DNA distortion and protein–DNA interactions (Sinha et al., 2023). Similarly, the level of chromatin occupancy by pioneer factors, influenced by motif content and local chromatin environment, has been shown to be a key determinant of their pioneering activity (Gibson et al., 2024). Given that crystallography studies show that IRF1 binding induces a ∼22° bending of the DNA (Escalante et al., 1998), clusters of IRF1 molecules may thereby cooperatively destabilize chromatin structure and promote remodeler recruitment.

A key finding of our study is that IRF1 cooperates with the SWI/SNF complex to unlock latent enhancers. While previous studies implicated IRF family members in SWI/SNF-mediated accessibility and activation of promoters (Ravi Sundar Jose Geetha et al., 2024; Ren et al., 2015), we show that IRF1 recruits BRG1-containing SWI/SNF to remodel chromatin at newly activated enhancers (C1–C3), both during IFNγ and LPS signaling. Pharmacological inhibition of BRG1 selectively dampened IRF1-driven chromatin accessibility and transcription, indicating that latent enhancers are particularly dependent on SWI/SNF activity (Chambers et al., 2023; Liao et al., 2024). Beyond recruitment, IRF1 also supports its pioneering capacity indirectly by inducing the expression of multiple chromatin regulators, including *Smarca4* (BRG1), *Kat2b*, *Kmt2c*, *Hdac9*, *Kat5*, and *Chd1*. This feed-forward mechanism likely reinforces broader chromatin accessibility and sustains enhancer activation beyond the initial wave of IRF1 binding.

This chromatin remodeling program extends beyond enhancer architecture to shape stress-adaptive gene modules in activated macrophages. Although BMDMs are largely non-proliferative, our data indicate that IRF1 engages regulatory elements near key checkpoint genes (*Rbl1*, *Hus1*, *Atm*), consistent with prior studies linking IFNγ signaling to cell-cycle arrest and apoptotic resistance that sustain antimicrobial responses (Armstrong et al., 2012; Kano et al., 1999; Xaus et al., 1999). Recent single-cell analyses on BMDMs indicate that macrophage inflammatory potential is strongly influenced by cell-cycle state, with G1-phase cells preferentially adopting pro-inflammatory programs (Daniel et al., 2023). By halting the G1/S transition, via p21 induction or cooperation with p53, IRF1 may stabilize a transcriptional state optimized for inflammatory and antimicrobial functions (Armstrong et al., 2012 ; Kano et al., 1999; Xaus et al., 1999). In parallel, IRF1 induces antioxidant and stress-response genes (*Gpx3*, *Glrx*, *Ptgs2*), buffering reactive oxygen species and promoting macrophage survival during peak effector activity (Mills et al., 2016). Together, these findings position IRF1 as a coordinator of cell-cycle related genes and stress resilience in activated macrophages.

In parallel to checkpoint and stress resilience, IRF1 also rewires core metabolism to meet the energetic and redox demands of activation. Immunometabolic adaptation is central to innate immune function (O’Neill & Pearce, 2016; Pearce & Pearce, 2013), yet the transcriptional drivers that connect cytokine signaling to these programs remain less clear, and our data suggest IRF1 is one such driver downstream of IFNγ. IRF1 deficiency markedly impairs induction of key glycolytic transporters and enzymes (*Slc2a1*, *Hk1, Hk3, Pfkp, Pkm, Ldha*) and prevents the IFNγ-induced shift from OXPHOS to aerobic glycolysis, consistent with the established IFNγ metabolic program (Wang et al., 2018). Chromatin and metabolite profiling further revealed that IRF1 directly couples transcriptional control to metabolic flux. IRF1 binds and opens an intragenic enhancer at the locus, promoting enhancer–promoter coupling and increased glycolytic capacity. Beyond glycolysis, IRF1 also occupies multiple TCA-related loci, including rapid binding at *Acod1* promoter and later engagement at *Sdhd*, coinciding with IRF1-dependent accumulation of TCA intermediates. Notably, IRF1-driven *Acod1* (IRG1) induction and itaconate accumulation link antimicrobial defense with redox and inflammatory regulation (Michelucci et al., 2013; Mills et al., 2018), while increased succinate and related metabolites are consistent with enhanced HIF-1α inflammatory signaling (Tannahill et al., 2013). Concurrently, IRF1 increases PPP engagement, supporting NADPH generation and glutathione-based redox buffering (Franchina et al., 2022; Nakamizo et al., 2023; Shah et al., 2024). This integrated transcriptional-metabolic program likely enables macrophages to tolerate oxidative stress while sustaining inflammatory outputs (Mills et al., 2016; Wang et al., 2018).

Finally, our findings demonstrate that IRF1 establishes long-lasting epigenetic imprints that persist beyond IFNγ withdrawal, characterized by stable H3K4me1 deposition and rapid reacquisition of H3K27ac upon restimulation. These “memory” features align with the concept of trained immunity, in which innate immune cells display heightened recall responses (Kaufmann et al., 2018; Moorlag et al., 2020; Netea et al., 2016). The IFNγ–IRF1 axis therefore represent a mechanistic route to epigenetic training, consistent with prior observations that type II IFN can override tolerance and reprogram myeloid responses (Chen & Ivashkiv, 2010; Kang et al., 2019). Furthermore, consistent with IFNγ being central to innate epigenetic memory, mycobacteria-induced trained immunity *in vivo* requires IFNγ, which imprints durable antimicrobial programming in macrophages following systemic BCG vaccination (Tran et al., 2024). In line with this, IRF1 is up-regulated in hematopoietic stem cells following BCG vaccination(Kaufmann et al., 2018) and IRF1 motif rank among the top enriched features at differentially accessible regions in β-glucan–trained neutrophils (Kalafati et al., 2020), although no mechanistic link was demonstrated at the time. This model may help explain earlier reports in humans of defective trained immunity in mononuclear cells derived from patients with STAT1-associated chronic mucocutaneous candidiasis but not STAT3-associated hyper-IgE syndrome, as STAT1 is required for IFNγ-driven IRF1 expression, while STAT3 does not comparably engage this axis (Ifrim et al., 2015). Collectively, these findings place IRF1 at the core of IFNγ-driven epigenetic training in innate immune cells.

As detailed above, these durable enhancers extend to metabolic loci (e.g., *Hk3*, *Hif1a*), supporting sustained glycolytic and respiratory remodeling during prolonged IFNγ exposure, a pattern that echoes trained immunity models in which a shift to glycolysis is fundamental for innate immune memory (Arts, Novakovic, et al., 2016; Keating et al., 2020; Otto et al., 2021), even if glycolytic reprogramming can be reversed under certain conditions (Wang et al., 2018). In parallel, IRF1-driven changes in TCA flux may reinforce epigenetic imprinting because intermediates such as fumarate, succinate, and acetyl-CoA act as cofactors for chromatin-modifying enzymes (Abhimanyu et al., 2022; Jiang et al., 2022; Noe & Mitchell, 2019; Sivanand et al., 2018; Tannahill et al., 2013). These results illustrate how IRF1’s metabolic imprint can synergize with chromatin modifications to shape trained immunity.

IFNγ signaling converges on pathways also implicated in BCG-induced trained immunity, where heightened glycolysis and TCA reorganization support stable histone modifications and enhanced immune responsiveness (Ferreira et al., 2024; Noe & Mitchell, 2019). Indeed, blocking these metabolic shifts abrogates immunological reprogramming (Abhimanyu et al., 2022; Arts, Carvalho, et al., 2016; Cheng et al., 2014; Keating et al., 2020; Otto et al., 2021), underscoring the deep functional interplay between metabolism and epigenetics. Consequently, while IRF1 KO macrophages may avoid some hyperinflammatory states (Berghout et al., 2013; Langlais et al., 2016), they also lose beneficial long-term memory capacity. Future research should clarify how IRF1 intersects with other signal-dependent TFs (e.g., NF-κB, STAT1, AP-1) to fine-tune epigenetic imprinting, thereby balancing protective immunity against pathological hyperactivation. Ultimately, our findings position IRF1 as a central driver that links epigenetic memory with metabolic adaptation, two essential processes that support epigenetic memory in macrophages.

## DATA SHARING STATEMENT

All original sequencing datasets will be available on NCBI Gene Expression Omnibus (GEO) under the following accession numbers: ATAC-seq (GSE317654), ChIP-seq (GSE317649), washout ChIP-seq experiments (GSE318866), SLAM-seq (GSE318866), and Hi-ChIP (GSE317657).

## AUTHORS CONTRIBUTIONS

JMA and DL designed the study. JMA performed most experiments, bioinformatic analyses, integrated multi-omics data, and drafted and revised the manuscript. RB managed the mouse colony, analyzed flow cytometry data, and contributed to writing and data presentation. MM conducted ATAC-seq experiments. AVIM and CFP performed cell respiration experiments. DL supervised all aspects of the project.

## ACKNOWLEDGMENTS

We thank HanChen Wang for his bioinformatics assistance, Hamed Najafabadi and Raquel Cuella Martin for technical assistance and constructive feedback. We thank Senthilkumar Kailasam at the Canadian Centre for Computational Genomics for support in processing the Hi-ChIP data. We are grateful to the Victor Phillip Dahdaleh Institute of Genomic Medicine for access to advanced facilities, including Illumina sequencing, and metabolomics (Nidia Lauzon). This work was supported by a joint Japan Agency for Medical Research and Development (AMED) and Canadian Institutes of Health Research (CIHR) Research Collaboration Grant to DL (#174122). DL is also supported by a Fonds de Recherche du Québec-Santé (FRQ-S) Chercheur-Boursier Junior 2 Award (doi.org/10.69777/348782). Equipment supporting this research were purchased with the concur of a Canada Foundation for Innovation John R. Evans Leaders Fund (CFI-JELF) grant to DL (#38033). CFP-Lab was supported by a Natural Sciences and Engineering Research Council of Canada (NSERC) Discovery Grant RGPIN-2024-04103 and by the Canada Foundation for Innovation, grant number 38858. CFP is the holder of the Canada Research Chair in Sustainable Integrative Parasitology (grant no. CRC-2024-00026). JMA was supported by the CONACYT I2T2 Northeastern Regional Scholarship, the Herbert A. Davis / M. Eve Cameron-Davis / Derek H. Davis Fellowship, and FRQ-S Doctoral Training Scholarship. MM was supported by a Postdoctoral Training Fellowship from Arthritis Society Canada and the Richard an Edith Strauss Foundation. RB received a Doctoral Training Scholarship from FRQ-S.

## METHODS

### Ethical statement

All mice were kept under specific pathogen-free conditions and handled according to the guidelines and regulations of the Canadian Council on Animal Care. Experimental protocols were approved by the McGill University Institutional Animal Care Committee (protocol number MCGL-8014).

### Bone marrow-derived macrophages

Bone marrow-derived macrophages (BMDMs) were differentiated from bone marrow cells isolated from the femurs and tibias of 8–12-week-old C57BL/6 (B6), termed wild-type (WT) or *Irf1^−/−^* female mice (Langlais et al., 2016). Briefly, bone marrow cells were cultured in high-glucose DMEM (Wisent, Cat. #319-005-CL), containing 10% heat-inactivated FBS (HI-FBS), Penicillin 100 U/mL + Streptomycin 100 µg/mL (Corning, 100× stock, Cat. #30-002-CI), and 20% L929 cell-conditioned medium (LCCM) as a source of macrophage colony-stimulating factor (M-CSF). On day 4, an additional 10% LCCM was added. On day 6, cells were harvested via a gentle wash with 1X PBS 5 mM EDTA. For each experiment, BMDMs were plated in DMEM with 10% HI-FBS, 20% LCCM, and antibiotics, and used the next day. Alternatively, cells were frozen slowly in 90% HI-FBS and 10% DMSO and stored at −80°C until further use.

### Flow cytometry

To assess the *in vitro* differentiation of BMDMs, cells were analyzed by flow cytometry (Ze5, Bio-Rad) viability using the Live/Dead Aqua Fixable Dead Cell Stain Kit (Thermo Fisher, Cat. #L34957) and for surface markers CD11b (eBioscience, 46-0112-80), F4/80 (eBioscience, 123114), and Ly6G (BioLegend, 128044). Live cells were gated based on Aqua dye exclusion, and successful macrophage differentiation was confirmed as >98% of live cells were F4/80⁺, Ly6G⁻, and CD11c⁻, confirming the absence of neutrophils and dendritic cells (Supplementary Figure 1A).

### Protein extraction and western blotting

BMDMs from WT and *Irf1⁻/⁻* mice were plated at 1×10⁷ cells per 10 cm tissue culture-treated dish and either left unstimulated or stimulated with 400 U/mL of murine IFNγ (R&D Systems, Cat. #485-MI-100) for 6 h. BMDMs cells were washed with ice cold PBS and lysed in RIPA lysis buffer (10 mM Tris-HCl, pH 8.0, 1mM EDTA, 0.5mM EGTA, 1% Triton X-100, 0.1% Sodium Deoxycholate, 0.1% SDS and 140mM NaCl) supplemented with a protease inhibitory cocktail (Aprotinin, Leupeptin and Pepstatin A at 0.1mg/ml each). Protein lysate concentrations were determined using the DC Protein Assay (Bio-Rad). 40 µg of proteins were separated on 10% stain-free polyacrylamide gels (Bio-Rad) and transferred onto PDVF membranes using the Trans-Blot Turbo system (Bio-Rad). Membranes were blocked in 5% skim milk in PBS Tween-20 0.1% and incubated with primary antibodies against IRF1 (R&D Systems, Cat. #AF4715, 1:200) or GAPDH (Cell Signaling Technologies, Cat. #2118) as loading control, followed by appropriate HRP-conjugated secondary antibodies. Signals were detected using Bio-Rad ECL substrate and imaged with the Vilber FX7 Gel Documentation System.

### ATAC-seq

One million WT and *Irf1^−/−^* BMDMs were plated in 6-well non tissue culture-treated plates and stimulated with IFNγ at 400 U/ml (R&D Systems, Cat. #485-MI-100) for different time points between 0-48 h. ATAC-seq was performed using the Omni-ATAC protocol (Buenrostro et al., 2013). Briefly, cells were washed with PBS and detached using PBS EDTA 5 mM. and the nuclei of 25,000 cells were isolated by incubating the cells on ice for 3 min in Lysis Buffer (Tween-20 0.1%, NP-40 0.1% and Digitonin 0.01%). Nuclei were incubated in Transposition buffer (TD Illumina buffer, 1x PBS, Tween-20 0.1%, Digitonin 0.01% and Tn5 Transposase) at 37°C for 30 min. DNA was isolated using Favorgen MicroElute Gel/PCR Purification Kit and used in PCR amplification for library generation and qPCR amplification to determine the number of total PCR cycles to perform. DNA was purified and library quality was assessed using Agilent High Sensitivity DNA Bioanalysis chip. Samples were sequenced on an Illumina NovaSeq 6000 in a paired-end 100 bp configuration.

### ATAC-seq analysis

Sequence quality was assessed using FastQC. Nextera adaptor sequences were removed using Trimmomatic 0.36 (Bolger et al., 2014) with the following parameters: ILLUMINACLIP:NexteraPE-PE.fa:2:30:10:2:keepBothReads. The resulting FASTQ files were mapped to the UCSC mouse mm10 reference genome using Bowtie 2.3.5 with the options: -k 1 --no-discordant --no-mixed (Langmead & Salzberg, 2012). Duplicate reads were marked using Picard (Institute, 2025) and removed using Samtools/1.10 (Li et al., 2009). Tag directories were generated with the makeTagDirectory function in HOMER (Heinz et al., 2010) and used to create normalized BigWig files at 1 bp resolution for visualization on the Integrated Genomics Viewer (IGV) (Robinson et al., 2011). Peak calling was performed using MACS2 version 2.1.1.20160309 with the parameters: -f BAMPE -q 0.01 --nomodel --shift -75 --extsize 150 (Zhang et al., 2008). Peaks were verified by retrieving the corresponding signal intensity from that sample’s and input tag directories. Fold change was calculated by comparing sample vs input DNA peak heights, and peaks were filtered based on height >5 count per million (CPM) and fold change >3. The significant peaks from all conditions were merged using HOMER’s mergePeaks function with the parameter -d 100, and the ATAC signal was retrieved for each condition using HOMER’s annotatePeaks.pl function with the option: -size 200. To identify kinetic patterns of ATAC-seq signal in WT and *Irf1-/-* BMDMs, log₂ mean-centering and K-means clustering were performed using Java Cluster 3.0. From the clustered peaks, a heatmap was generated by annotating the normalized tag read distribution within ±2 kb of the peak center using HOMER’s annotatePeaks.pl function (-size 4000 -hist 50 -ghist) and visualized using Java TreeView. To analyze peak height kinetics, we identified the maximum value across all conditions for each peak and normalized heights relative to this maximum. Kinetic plots were generated by plotting these normalized average peak heights across all time points to visualize temporal changes.

### ATAC-seq footprinting analysis

Differential transcription factor (TF) binding analysis was performed using the TOBIAS software (Bentsen et al., 2020). All 879 core vertebrate motifs from the JASPAR database were compiled using the FormatMotifs function. ATAC-seq signals from WT and *Irf1-/-* datasets were corrected for Tn5 insertion bias using the ATACorrect and used for differential binding analysis between time points and conditions using the BINDetect function. Multiple testing correction was done using Bonferroni adjustments in R, and results were visualized as volcano plots. For clustering of TF binding kinetic, we identified the top 100 motifs exhibiting the most variability across time points in WT BMDMs. Footprinting scores for these motifs were re-annotated using the FootprintScores function to display binding intensity over time. Binding scores were log₂ mean-centered, and K-means 4 clustering was performed using Java Cluster 3.0 and visualized with Java TreeView. IRF1 footprinting in cluster 1 was visualized using heatmaps and aggregate plots generated with TOBIAS PlotHeatmap and PlotAggregate functions, respectively.

### Motif analysis

*De novo* motif analysis was performed on all clusters using HOMER’s findMotifsGenome.pl function with a size parameter of 100 bp (Heinz et al., 2010). Motifs with a p-value less than 1e-11 were selected and redundancies were filtered out. For each cluster, motif co-occurrence matrices were generated using their corresponding position weight matrices (PWMs) and annotated with annotatePeaks.pl using the - m function to query for a curated set of 17 PWMs. The motif occurrence data were transformed into binary matrices and converted to numeric values. Co-occupancy percentages and co-association matrices for transcription factor pairs were calculated. An igraph object was created for visualization using the ggraph package in R, with nodes sized according to peak percentages; motifs below a presence of 5% were filtered out to produce the final network graph. A *de novo* motif extension analysis was carried out using findMotifsGenome.pl with parameters -size 100 -len 34 to identify IRF1 motif arrays. To assess IRF1 motif distribution within peaks, ATAC-seq peaks were first centered on the IRF1 motif using annotatePeaks.pl with the -size 200 -center option, and then annotated for enrichment of the IRF1 motif using annotatePeaks.pl (-size 1000 -hist 10 -m), and results were visualized in R using ggplot2. Motif count categories were established by querying for the IRF1 motif across all identified ATAC peaks using annotatePeaks.pl -m; peaks were binned in R based on the number of IRF1 motifs per peak, and cluster participation in each category was determined. A heatmap of tag read distribution surrounding ±1 kb of the ATAC peak center was generated by annotating IRF1 ChIP-seq signal at 3 h post-IFNγ stimulation in WT BMDMs using annotatePeaks.pl with parameters -size 2000 -hist 50 and visualized in Java TreeView compared to input.

### ChIP sequencing and quantitative PCR (qPCR)

20 million BMDMs from WT and *Irf1⁻/⁻* mice were plated in 15 cm dishes, and the next day were treated or not with 400 U/ml IFNγ (R&D Systems, 485-MI-100/CF) for different duration; all conditions were ended simultaneously. ChIPs were performed as previously described (Langlais *et al.,* 2016) with few modifications. Briefly, BMDMs were crosslinked for 10 min at room temperature with 1% formaldehyde added in the culture medium, after which the medium was removed and crosslink was stopped with ice-cold PBS containing 0.125 M glycine for 5 min. Nuclei were prepared by sequential incubation on ice for 5 min in buffer A (10 mM Tris-HCl pH 8, 10 mM EDTA, and 0.25% Triton X-100) and for 30 min in buffer B (10 mM Tris-HCl pH 8, 1 mM EDTA, and 200 mM NaCl; all buffers included protease inhibitors). Nuclei were resuspended in a sonication buffer (10 mM Tris-HCl pH 8, 1 mM EDTA, 0.5% SDS, 0.5% Triton X-100, 0.05% NaDOC, and 140 mM NaCl) and fragmented to a size range of 100-500 bp using a digital sonifier (Branson Ultrasonics) at 80% amplitude, 7 minutes, ON / OFF 30 seconds. Sonicated chromatin was diluted 1:3 in ChIP dilution buffer and incubated overnight on a rotating platform at 4°C with a mixture of 40 µL protein G Dynabeads (Invitrogen) prebound with 3 µg of IRF1 (R&D, AF4715), 3 µg BRG1 (Abcam, ab10641), 3 µg H3 (Abcam, ab1791), 3 µg H3K4me1 (Abcam, ab8895), 6 µg H3K27ac (Abcam, ab4729), 1 µg H3K27me3 (CST, 9733) and 7 µg H3K9me2 (Abcam, ab1220). Immune complexes were washed sequentially for 2 min at room temperature with 1 mL of the following buffers: wash B (1% Triton X-100, 0.1% SDS, 150 mM NaCl, 2 mM EDTA, 20 mM Tris-HCl pH 8), wash C (1% Triton X-100, 0.1% SDS, 500 mM NaCl, 2 mM EDTA, 20 mM Tris-HCl pH 8), wash D (1% NP-40, 250 mM LiCl, 1 mM EDTA, 10 mM Tris-HCl pH 8), and TEN buffer (50 mM NaCl, 10 mM Tris-HCl pH 8, 1 mM EDTA). After de–crosslinking by overnight incubation at 65°C, the DNA treated with RNase A and Proteinase K, followed by purification on QIAquick PCR purification columns (Qiagen). ChIP efficiency was assessed by qPCR relative to the input DNA diluted to 10 pg/ul for TFs, and relative to H3 levels for histone mark using the 2X Universal SYBR Green Fast qPCR Mix (ABclonal) for known enrichment sites using primers listed in Table S1. The ChIP-seq libraries were prepared using the Kapa Hyperprep ChIP library kit (Roche Molecular Systems) and sequenced on a S4 flowcell on a NovaSeq 6000 sequencer in a paired-end 100-bp configuration.

### ChIP-seq analysis

Sequence quality assessment, mapping to the mm10 reference genome and processing was performed as described for ATAC-seq. TF peak calling was performed using MACS2 version 2.1.1.20160309 with the parameters -f BAMPE -q 0.01. The following published BMDM ChIP-seq datasets were retrieved from the Gene Expression Omnibus (GEO) database and reanalyzed using the parameters described above: H3K4me1 at baseline (0 h), after IFNγ stimulation (4 h), and under washout conditions (24 h IFNγ followed by a 48 h rest), as well as H3K4me3 at 0 h and 3 h IFNγ (Ostuni et al., 2013), and PU.1 and IRF8 at 0, 0.5, and 3 h of IFNγ stimulation (Langlais et al., 2016).Heatmaps were generated by annotating the normalized tag read distribution within ±1 kb (for TFs) or ±2 kb (for histones) of the ATAC-seq determined peak center using HOMER’s annotatePeaks.pl function (-size 4000 -hist 50 - ghist). The resulting data were visualized using Java TreeView. Density plots of tag read distribution for each clusters of peaks were generated using annotatePeaks.pl with -size 4000 -hist 10 and visualized in R using ggplot2 for IRF1, ATAC-seq, H3K27ac, H3K27me3, and H3K9me2 signals. To determine the kinetic of IRF1 binding, chromatin accessibility (ATAC) and histone modifications (H3K27ac, H3K9me2, H3K27me3) amongst peaks clusters 1-3, normalized signal was retrieved using annotatePeaks.pl with the option -size 200 and the resulting data were scaled between 0-1. Time series for each mark were plotted, and enrichment analysis was performed, and halftime of change was determined by calculating the time point at which the scaled signal reached 50%.

For genome-wide IRF1 binding analysis, the identified IRF1 peaks were merged across time points using HOMER’s mergePeaks function with the parameter -d 100 and filtered based on fold change over input (>5) and height>4 CPM. The peaks were then ranked by IRF1 signal intensity and divided into four binding classes based on signal strength (weak, moderate, strong, and very strong). Then, heatmaps of IRF1 and ATAC signals for each class were generated in R using the ggplot package.

IRF1 binding peaks were analyzed for proximity to genes within key metabolic pathways (Figure 6A) using the GenomicRanges and GenomicFeatures packages in R (Lawrence et al., 2013). Gene annotations were retrieved from the “TxDb.Mmusculus. UCSC.mm10.knownGene” database, and gene regions were extended by ±2 kb upstream and downstream. Overlaps between extended gene regions and IRF1 peaks were calculated. Visualization was performed using ggplot2 to display the proportion of genes with IRF1 peaks and the average number of peaks per gene for each pathway.

### Training experiment and analysis

For the washout and resting experiments (Figure 8), BMDMs were subjected to four stimulation regimes: untreated and cultured for 7 days; 24 h of IFNγ stimulation followed by washout and a 6-day of rest; 4 days of untreated culture followed by IFNγ stimulation for 24 h and then washout and 2-day rest; or IFNγ stimulation for 24 h followed by washout and 6-day rest. At endpoint, cells were harvested for ChIP-seq analysis of IRF1 and histone marks (H3K4me1, H3K27ac, H3K9me2).

Aggregate H3K4me1, H3K27ac, and H3K9me2 signals were quantified at C1-3 enhancers in naive, IFNγ-trained, and restimulated BMDMs. Differential H3K4me1 enrichment was also assessed at C1-3 ATAC-seq peaks using HOMER’s annotatePeaks.pl function with the option -size 200; CPM and log₂ fold changes between conditions (resting versus washout) were computed and a volcano plot was generated using ggplot2 to visualize differential enrichment.

### SLAM-seq

BMDMs from WT and *Irf1⁻/⁻* mice were generated from three independent mice per genotype (biological replicates) and plated at a density of 500,000 cells per well in 12-well tissue culture-treated plates containing 2 mL of LCCM containing complete media. The following day, cells were stimulated with 400 U/mL IFNγ (R&D Systems, Cat. #485-MI-100) for either 0, 1, 3, 6, 12, or 48 h. Thirty minutes before harvest, 4-thiouridine (4SUTP) (Jena Bioscience, Cat. #NU-1156L) was added to each well at a final concentration of 200 µM for metabolic labeling, as previously described (Herzog et al., 2017). After 30 minutes, media was removed, cells washed with ice cold PBS and lysed with 1 mL of Qiazol. RNA was extracted, treated with iodoacetamide (Biobasic, Cat. #IB0539) and purified using the Monarch Total RNA Mini Prep Kit (New England Biolabs, Cat. #T2010S) before quantification via BioDrop (Biochrom). Samples were sequenced on an Illumina NovaSeq 6000 in paired-end 100 bp obtaining an average 50 million reads per sample.

### SLAM-seq analysis

Sequence reads quality was controlled using FastQC v0.12.0 and trimmed with Trimmomatic v0.39 using default parameters. Trimmed reads were mapped for both total and nascent RNA fractions: using Hisat2 (Kim et al., 2015) with the option -k 1 for total RNA, and Hisat3n (Zhang et al., 2021) with options --base-change T,C and --no-repeat-index for nascent RNA. Non-nascent reads were filtered from the resulting Hisat3n BAM files using Samtools (Li et al., 2009) by selecting on the "Yf:i:" tag. Sorted BAM files were generated using Samtools, and read counts were obtained with FeatureCounts (Liao et al., 2014) from the SUBREAD v2.0.3 package using parameters -p -t exon -g gene_id. Gene expression converted to counts per million (CPM) and expressed as log2-fold changes relative to the group average of their respective 0 h time points. Genes with expression levels above a minimal cutoff (mean CPM > 20) were retained for downstream analyses. For BigWig file generation, a scaling factor was calculated by dividing 1 by the number of reads mapped to exons (excluding ribosomal RNA reads) and normalizing to one million reads. Scaled BedGraph files were generated using genomeCoverageBed, and BedGraph files were converted to BigWig format using the UCSC wigToBigWig tool (Kent et al., 2010). Principal component analysis (PCA) was performed on CPM values to examine variance in both total and nascent RNA reads, and visualized using ggplot2 in R. For nearest gene-to-cluster expression analysis, we identified the nearest genes to each peak with annotatePeaks.pl and filtered based on an absolute distance cutoff of 10 kb from transcriptional start sites, focusing on the single nearest gene.

Gene Ontology (GO) enrichment analysis was performed using the enrichGO() function of clusterProfiler R package (Yu et al., 2012). Gene lists were defined as genes located within ±50 kb of transcription start sites and gene symbols were converted to Entrez IDs using the bitr() function.

Enrichment of GO biological process terms was assessed using Benjamini–Hochberg multiple-testing correction with a q-value cutoff of 0.05. Fold enrichment for each GO term was calculated from observed and background gene ratios, log₂-transformed, and the top six terms per gene list were selected based on adjusted p-values. These enriched GO terms were visualized in a heatmap to summarize functional differences across clusters. Separately, expression heatmaps were generated to visualize log₂-fold change mean-centered gene expression over time. For cluster 1, redundant GO terms were removed by selecting those with the highest p-values, resulting in four GO terms. Gene lists corresponding to these GO terms were used to extract expression data from the nascent RNA fraction, which were then visualized using ggplot2. Similarly, a gene expression heatmap was created for specific metabolic pathways of interest, with genes ordered according to their positions in the pathway (Figure S6). A distinct heatmap of chromatin remodeler expression patterns was generated using the GOBP_CHROMATIN_REMODELING gene set (GO:0006338). For pioneering and metabolic gene panels, individual gene expression analysis was performed by calculating the mean and standard deviation of gene expression levels. Statistical significance between WT and knockout (KO) conditions across time was assessed using a two-way ANOVA (genotype × time), followed by post-hoc pairwise comparisons at each time point. Expression profiles of pathways were visualized using ggplot2, with individual panels for each gene. Gene set enrichment analysis was performed using the clusterProfiler package (Yu et al., 2012) on pre-ranked gene lists ordered by signed −log10(P value) from differential expression statistics, with enrichment evaluated against curated MSigDB gene sets (Reactome, Gene Ontology, and MH collections) using permutation-based testing, normalized enrichment scores, and multiple-testing correction (Subramanian et al., 2005).

RNA-seq data from macrophages treated or not with BRM014 (1 μM), a BRM/BRG1 ATPase inhibitor, in the context of stimulation with Lipid A for 4 hours (GEO GSE242803) (Liao et al., 2024) were downloaded and processed using the pipeline described above. Differential gene expression analysis was performed at the Lipid A 4h time point comparing the BRM014 treated and non-treated groups, and the log2 odds ratio of downregulated genes was calculated for each cluster to assess the impact of BRG1 inhibition on IRF1-dependent gene regulation.

### Hi-ChIP

20 million BMDMs from WT and *Irf1⁻/⁻* mice were plated in 15 cm tissue culture-treated dishes as described above and stimulated with IFNγ (400 U/mL, R&D Systems) for 0 – 48 h. Crosslinking and chromatin immunoprecipitation were performed following the Dovetail Genomics Hi-ChIP protocol using the IRF1 antibody (R&D Systems, AF4715). Libraries were prepared with Dovetail i7 and i5 dual index primers and sequenced on an Illumina NovaSeq 6000 generating 100 bp paired-end reads. Data preprocessing was conducted using FitHiChIP v11.0 (Bhattacharyya et al., 2019) with a FDR cutoff of 0.1. Loop calling utilized the merged list of IRF1 peaks from all conditions, applying stringent filtering criteria of >6 fold enrichment over input and >6 CPM. Resulting BEDPE files were visualized in IGV, highlighting key genes with IRF1-mediated chromatin looping. To visualize chromatin interactions between clusters, Hi-ChIP interaction anchors were assigned to their nearest ATAC-seq cluster. Interactions with unassigned anchors were excluded, and the number of interactions between each cluster pair was quantified. Inter-cluster interactions were visualized as arcs using ggraph and igraph in R, with arc widths proportional to interaction frequency. We also assessed the relationship between the number of IRF1 motifs (0 to 23) within ±100 bp of ATAC-seq peaks and chromatin loop participation. Genomic regions containing IRF1 motifs were converted into GRanges objects using the GenomicRanges package (Lawrence et al., 2013). Hi-ChIP loop anchors were expanded by 2 kb upstream and downstream and overlaps between motif regions and expanded anchors were identified. Participation proportions for each motif count were calculated based on these overlaps. Logistic regression analysis using the glm function in R was applied to model the relationship between motif count and loop participation likelihood.

### OCR measurements

OCR measurements were conducted using the Resipher platform (Lucid Scientific), which allows real-time continuous monitoring of oxygen consumption in cellular metabolism. WT and *Irf1⁻/⁻* BMDMs were plated at a density of 50,000 cells per well in a 96-well plate, using four biological replicates per condition. After allowing the cells to rest for 24 hours to promote adhesion, cells were treated with IFNγ (400 U/mL), LPS (100 ng/mL), or β-glucan (5 μg/mL; (1→3)-β-D-glucan from *Alcaligenes faecalis*), and oxygen consumption was measured continuously for five days in a 5% CO₂ incubator at 37 °C. Data were reported in units of femtomoles per square millimeter per second (fmol/mm²/s).

### Metabolomics

BMDMs from WT and *Irf1⁻/⁻* mice were seeded at 5 million per 10-cm tissue-culture dish and allowed to adhere for 24 h. Cells were then stimulated with IFNγ 400 U/mL for 0–48 h; all time points were harvested concurrently. The culture medium was aspirated and monolayers washed twice with room-temperature PBS. Metabolite extraction was initiated by adding 300 µL of pre-chilled (−20 °C) 3:1 (v/v) 1-butanol:methanol (MilliporeSigma) directly to each dish; plates were kept on ice while cells were scraped and the suspension transferred to 1.5-mL microcentrifuge tubes on ice. Samples were dried by vacuum evaporation at 25 °C for 3 h. Dried extracts were resuspended in 40 µL LC-MS grade water and 200 µL internal-standard methanol (LC-MS grade; VWR) containing stable isotope labeled standards (fructose-13C6, methylmalonic acid-d3; Cayman Chemical) and mixed by vortexing. Samples were shaken at 1,200 rpm for 10 min at 25 °C and centrifuged at 16,000 × g for 10 min at 4 °C. 120 µL of the resulting supernatant were transferred to fresh tubes for biphasic extraction by adding 400 µL methyl tert-butyl ether (MTBE; VWR) and 110 µL LC-MS grade water, followed by vortexing for 15 s and centrifugation at 2,000 × g for 10 min at 25 °C. The lower aqueous phase, containing polar metabolites, was collected and split for LC-MS/MS and GC-MS/MS workflows.

For targeted LC-MS/MS, 90 µL of the aqueous phase were dried by vacuum evaporation at 25 °C for 3 h and reconstituted in 26 µL 0.1% formic acid in water (LC-MS grade; Fisher Scientific and VWR). Reconstituted samples were vortexed, centrifuged to remove particulates, transferred to LC autosampler vials and analyzed on a Shimadzu LCMS-8060NX system. For targeted GC-MS/MS, 90 µL of the aqueous phase were dried in amber autosampler vials and derivatized by adding 40 µL methoxyamine hydrochloride in pyridine (20 mg/mL; MeOX and pyridine, MilliporeSigma and VWR) and shaking at 37 °C for 30 min at 750 rpm. Twenty microliters of N-methyl-N-trimethylsilyl-trifluoroacetamide (MSTFA; MilliporeSigma) were then added and samples were shaken for an additional 30 min at 37 °C, centrifuged at 5,000 × g for 5 min at 25 °C, and injected on a Shimadzu GCMS-8040NX system.

### Metabolomic analysis

Metabolite signal intensities were normalized to internal standards prior to statistical analysis. Downstream processing was performed in R using custom scripts. Intensity matrices from GC–MS and LC–MS were imported and inspected for completeness; missing values were imputed using a sample-wise minimum-intensity approach to ensure retention of low-abundance compounds. Intensities were normalized by median scaling across samples to correct for global differences in total ion abundance, then log₂-transformed with a small offset to stabilize low-abundance measurements. Metabolites that were constant across all samples after normalization (zero variance) were excluded, while low-abundance but variable compounds were retained. Data quality was evaluated through principal component analysis (PCA), computed on mean-centered values using the prcomp function and generated with ggplot2. Compound-level summaries were obtained by calculating the mean and standard deviation across replicates for each genotype and time point. Differential abundance analysis was carried out using the limma package (Ritchie et al., 2015). For the GC–MS dataset, a linear model was fitted for each metabolite followed by empirical Bayes moderation with the eBayes function (Smyth, 2004). Two contrasts were tested: (i) WT 0 h versus WT 48 h and (ii) KO versus WT at 24 h. Moderated statistics were used to derive log₂-fold changes and adjusted p-values using Benjamini–Hochberg correction. Volcano plots were generated from these results using ggplot2, with metabolites plotted by log₂-fold change and −log₁₀ adjusted p-value, with a significance threshold established at adjusted p < 0.05. For selected metabolites of interest, genotype-specific differences at individual time points were assessed using two-sided Student’s t-tests. Lastly, the GSH/GSSG ratio was computed for each sample by dividing normalized GSH intensities by GSSG intensities, then summarized by genotype and time point and plotted as mean trajectories with shaded standard deviation using ggplot2.

