## Supplementary Figures and Tables for "Pioneer factor IRF1 unlocks latent enhancers to rewire chromatin and immunometabolism in inflammatory macrophages"

*Pioneer factor IRF1 unlocks latent enhancers to rewire chromatin and immunometabolism in inflammatory macrophages via SWI/SNF-dependent remodeling*

Jose-Mauricio Ayala [1, 2], Rebecca Bellworthy [1 *], Mathieu Mancini [2, 4 *], Ana Victoria Ibarra-Meneses [3], Christopher Fernandez-Prada [3], David Langlais [1, 2, 4]

[1] Department of Human Genetics, Victor Phillip Dahdaleh Institute of Genomic Medicine, McGill University, Montreal, QC, Canada, H3A 0G1

[2] McGill University Research Centre on Complex Traits, Montreal, QC, Canada

[3] Département de Pathologie et Microbiologie, Faculté de Médecine Vétérinaire, Université de Montréal, Saint-Hyacinthe, Québec, Canada, J2S 2M2

[4] Department of Microbiology and Immunology, McGill University, Montreal, QC, Canada, H3A 2B4

[*] Equal contributions

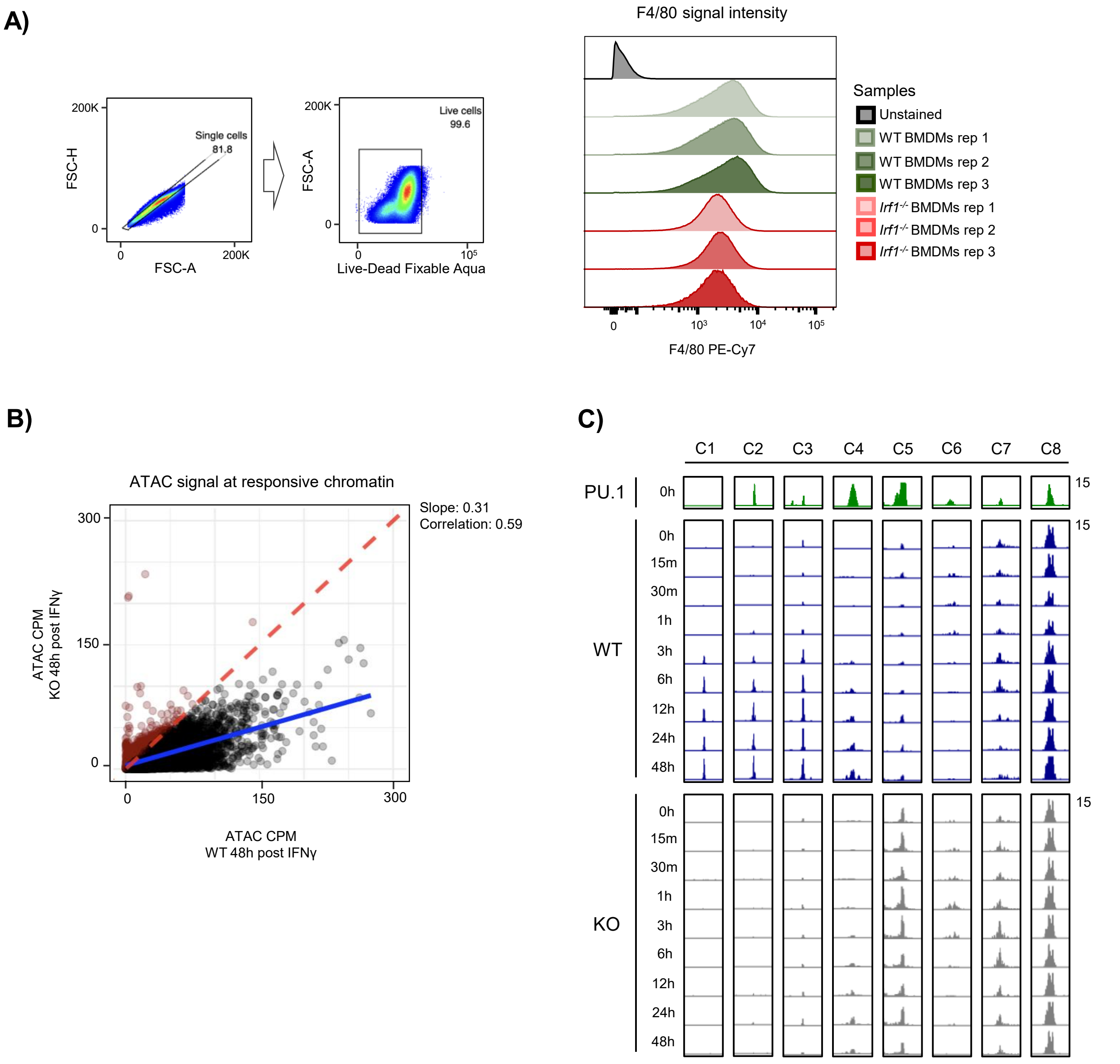


**Supplementary Figure 1. IRF1 influences IFNγ-induced chromatin accessibility without altering core macrophage identity, related to Figure 1**

**(A)** Representative flow cytometry plots showing the gating strategy for singlets (FSC-H vs FSC-A), live/dead discrimination (left), and BMDM markers. Histograms of F4/80 expression (right) in one batch of 3 WT (green) and 3 IRF1 KO (red) BMDMs. **(B)** Scatter plot comparing normalized ATAC-seq counts at 48 h post–IFNγ in WT (x-axis) and IRF1 KO (y-axis) BMDMs (points = genomic loci); red points = loci higher in KO, black = loci higher in WT; fitted slope = 0.31 (blue line). **(C)** Genome browser snapshots of selected example loci (columns) for each cluster; rows display normalized PU.1 ChIP-seq signal (green) and normalized ATAC-seq signal (WT = blue, KO = grey).

A)

*De novo* motif analysis of cluster 1

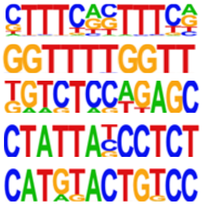

| Most similar known TF motif | Motif | P-value | % of targets |
|---|---|---|---|
| IRF1 | | $1 \times 10^{-515}$ | 60.55% |
| IRF5 | | $1 \times 10^{-191}$ | 13.35% |
| ZBTB26 | | $1 \times 10^{-30}$ | 1.12% |
| Spz1 | | $1 \times 10^{-22}$ | 0.86% |
| ZBTB32 | | $1 \times 10^{-22}$ | 0.86% |

B)

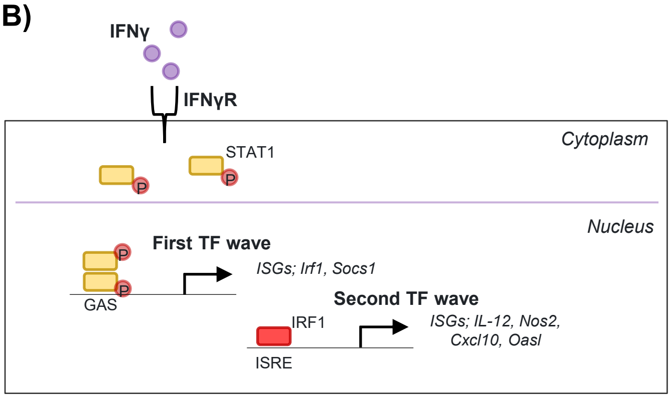


**Supplementary Figure 2. IRF1 is the dominant transcription factor at pioneered enhancers, related to Figure 2.**
**(A)** Table of *de novo* motifs enriched in cluster 1 peaks, with the most similar TF motif, p-value and percentage of sites bearing each motif. **(B)** Schematic in WT BMDMs illustrating three successive waves of transcription factor motif occupancy derived from TOBIAS clusters (Figure 2C).

### Supplementary Figure 3

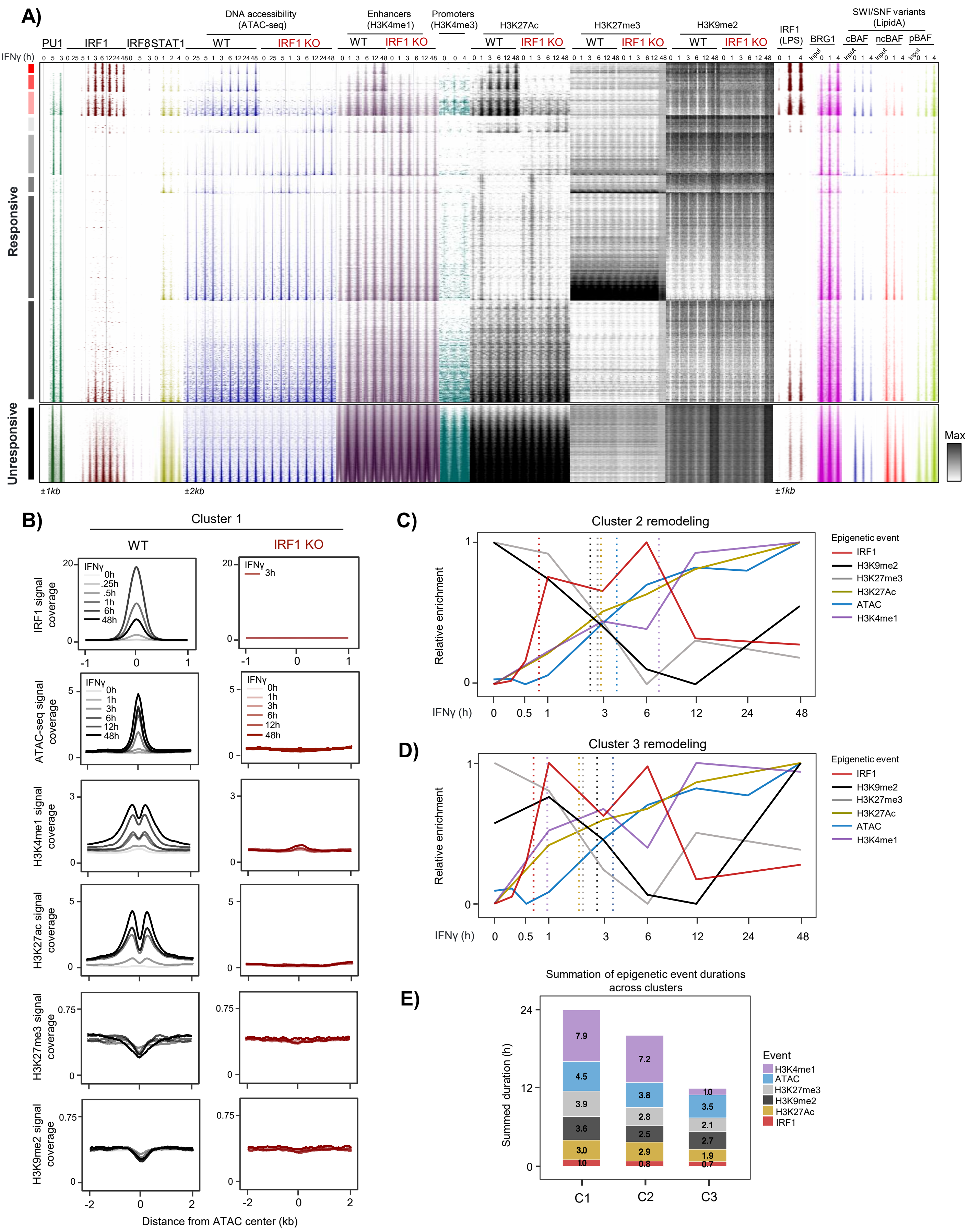

**Supplementary Figure 3. IRF1-dependent chromatin remodeling in macrophages, related to Figure 3.**

**(A)** Multi-omic heatmaps showing ChIP–seq signals (PU.1, IRF1, IRF8, STAT1 and SWI/SNF components), normalized ATAC-seq accessibility and histone modification tracks (H3K4me1, H3K4me3, H3K27ac, H3K27me3, H3K9me2) across matched time points (0–48 h); rows = regions, columns = time points. **(B)** Density plots of aggregated IRF1, ATAC-seq, H3K4me1, H3K27ac, H3K27me3 and H3K9me2 across matched time points for WT and IRF1 KO BMDMs. **(C, D)** Line graphs of timelines depicting sequential events at Cluster 2 and 3 sites. The half-time ($t_{1/2}$) to reach 50% of each signal's maximum was used to order temporal steps; indicated by a colored dashed line. **(E)** Stacked bar plots of cumulative $t_{1/2}$ (hours) for IRF1 binding (red), H3K4me1 (purple), chromatin opening (ATAC, blue), H3K27ac (yellow) and removal of repressive marks (H3K27me3 light gray; H3K9me2 dark gray) across Clusters 1–3; numeric labels indicate the number of hours per step.

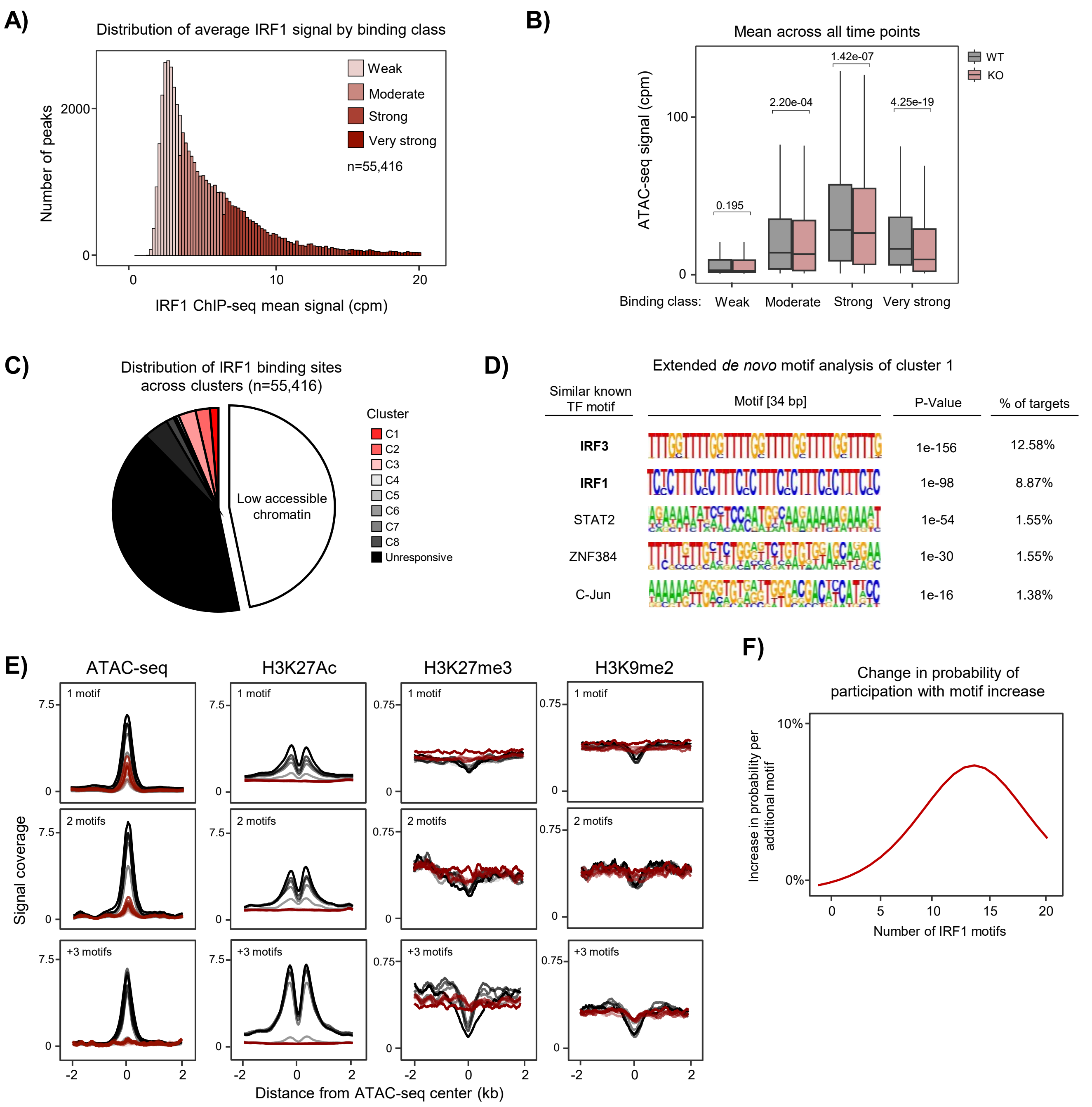


**Supplementary Figure 4. IRF1 binding strength and motif number determine chromatin accessibility and histone modifications, related to Figure 4.**

**(A)** Histogram of the number of IRF1 ChIP-seq peaks (n = 55,416) stratified into weak, moderate, strong and very-strong binding categories based on normalized ChIP-seq signal intensity. **(B)** Boxplots of normalized ATAC-seq accessibility grouped by IRF1 binding strength (weak → very strong). WT vs IRF1 comparison by Wilcoxon test. **(C)** Pie chart of the proportion of IRF1 binding categories across ATAC-seq clusters; ~57% of sites are in low-accessibility clusters. **(D)** HOMER motif-enrichment analysis of extended 34-bp de novo motifs at ATAC-seq Cluster 1 showing enrichment for tandem IRF motifs. The table lists the most similar known motif, motif sequence, p-value and percent sites. **(E)** Aggregate signal graphs in WT and IRF1 KO BMDMs of normalized ATAC-seq, H3K27ac, H3K27me3 and H3K9me2 stratified by IRF1 motif categories. **(F)** Plot of the fraction of IRF1-bound sites forming Hi-ChIP loops versus motif count with a fitted trend line; an increase peaking near ~15 motifs is shown.

### Supplementary Figure 5

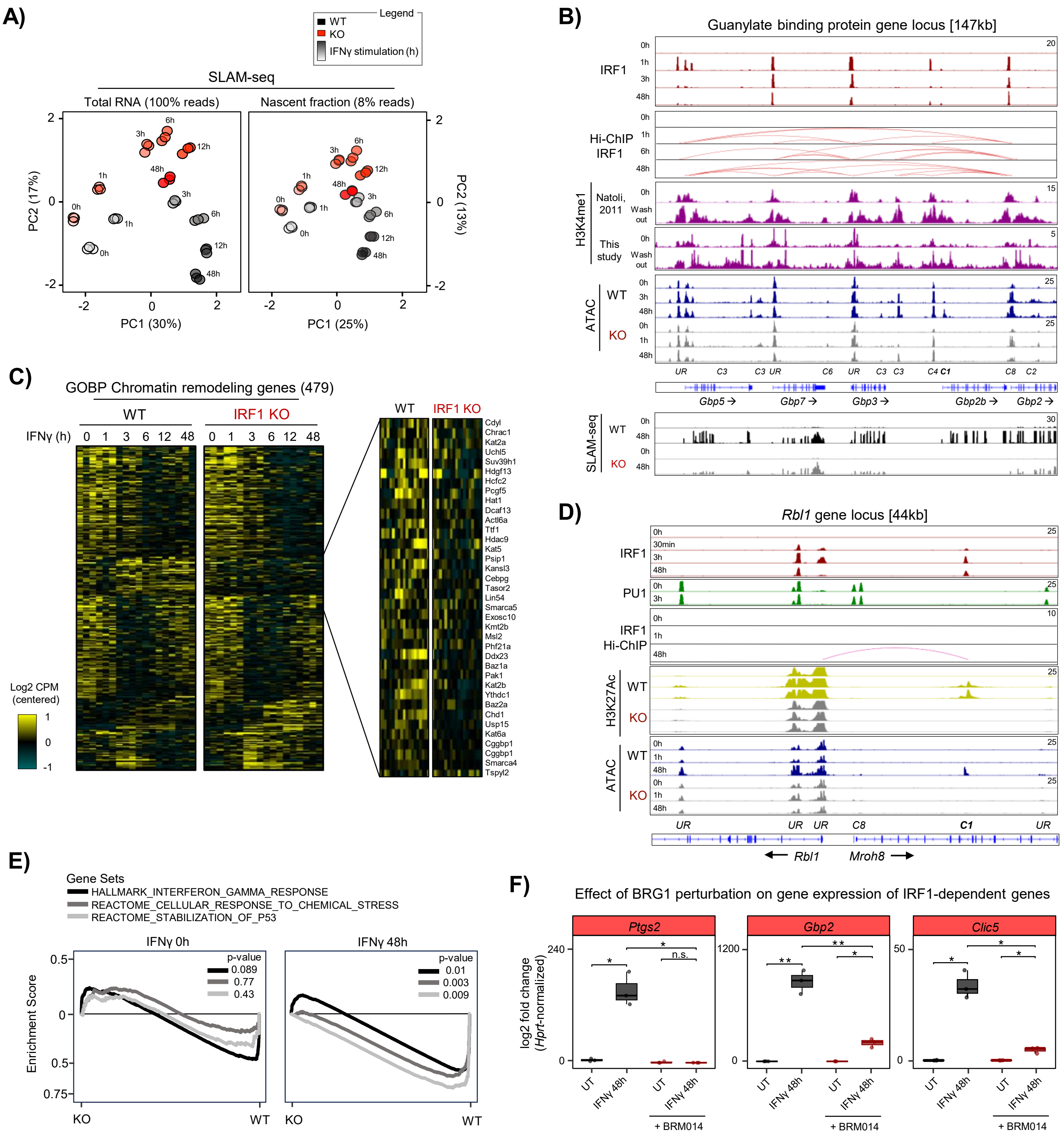

**Supplementary Figure 5. IRF1 controls chromatin remodeling factors through the action of SWI/SNF, related to Figure 5.**

**(A)** Principal-component analysis (PCA) of total RNA and nascent RNA fractions from WT and IRF1 KO BMDMs collected across matched IFNγ time points. Each point represents a sample.; PCA was performed separately for total and nascent RNA datasets. [Total fraction = Hisat2 mapping, Nascent fraction = Hisat3N mapping; n = 3/group]. **(B)** Genome browser tracks at the *Gbp* locus showing IRF1 ChIP-seq, Hi-ChIP (IRF1) interactions, H3K4me1 profiles, ATAC-seq (WT and IRF1 KO), and SLAM-seq signal (WT and IRF1 KO). Tracks are shown across stimulation and washout conditions. [UR = Unresponsive]. **(C)** Heatmap of nascent RNA expression for genes listed in the GOBP_CHROMATIN_REMODELING category [GO:0006338], in WT and IRF1 KO BMDMs following IFNγ stimulation. Rows represent genes and columns represent time points and genotypes; genes are ordered by unsupervised hierarchical clustering. **(D)** Genome browser tracks at the *Rbl1* locus showing IRF1, PU.1 and H3K27ac ChIP–seq, Hi-ChIP interactions and ATAC–seq in WT and IRF1 KO BMDMs. **(E)** Gene Set Enrichment Analysis (GSEA) plots for p53 stabilization, interferon gamma response, and cellular response to chemical stress gene sets in WT and IRF1 KO BMDMs at 0 h and 48 h IFNγ. **(F)** Boxplots showing counts-per-million (CPM) expression of *Ptgs2* (Cluster 1), *Gbp2* (Cluster 2), and *Clic5* (Cluster 3) in BMDMs untreated or stimulated with IFNγ for 48 h, in the presence or absence of BRM014. Individual points indicate biological replicates; boxes represent the interquartile range with the median; p-values are indicated (*$p < 0.05$, **$p < 0.01$; n.s.).

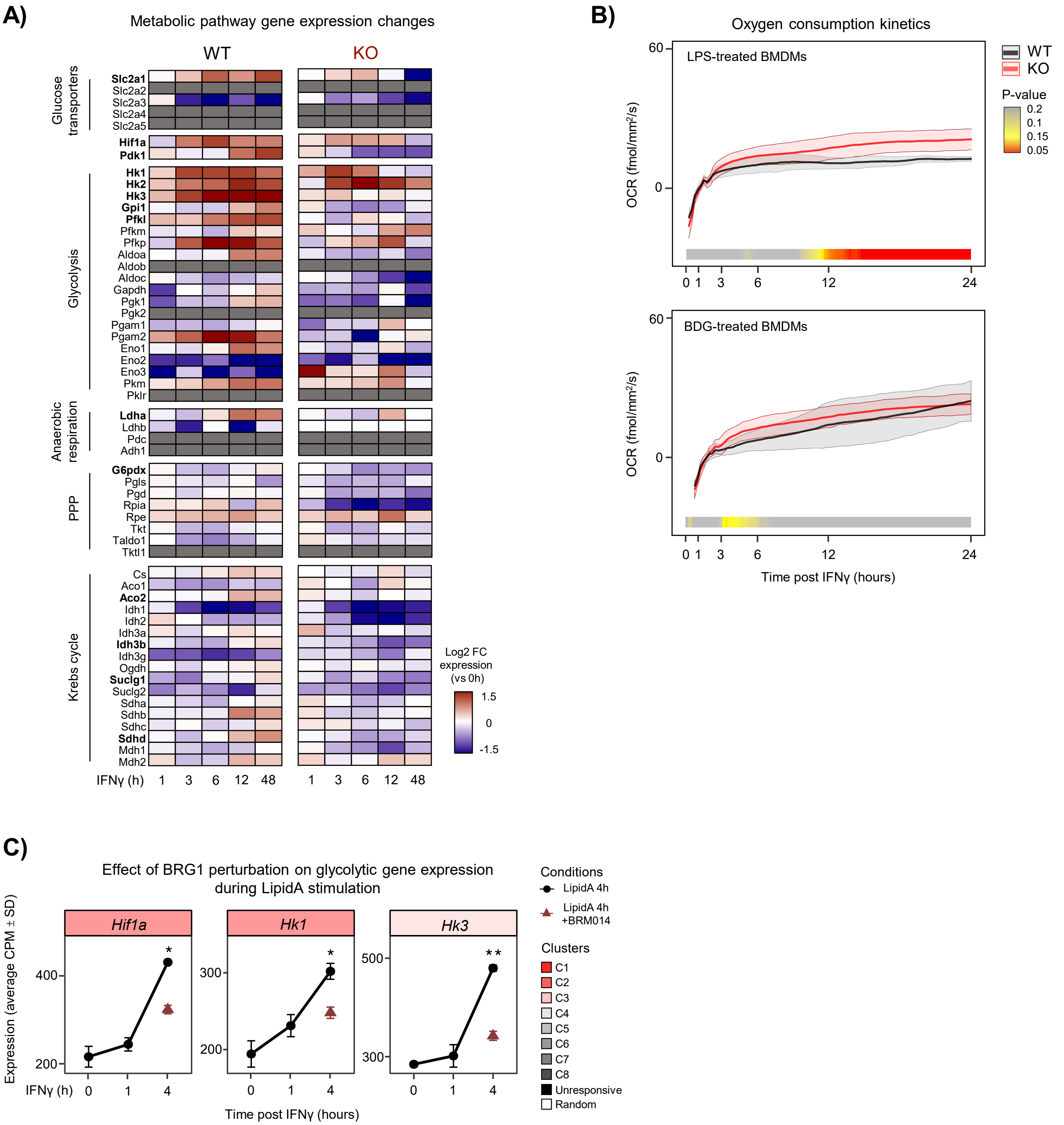


**Supplementary Figure 6. IRF1 coordinates metabolic reprogramming in macrophages, related to Figure 6.**
**(A)** Heatmap of metabolic pathway gene expression showing log2 fold-change relative to 0 h (range +1.5 to −1.5) in WT and IRF1 KO BMDMs; rows correspond to genes grouped by metabolic pathway and columns to IFNγ time points (1, 3, 6, 12, 48 h) and genotype. **(B)** Ribbon plots showing oxygen consumption rate (OCR) kinetics (mean ± SD, n = 4 per condition) for WT and IRF1 KO BMDMs following LPS treatment (top) or BDG treatment (bottom). Heat maps below each plot display rolling p-values from T-tests for WT versus KO comparisons, with red indicating $p < 0.05$. **(C)** RNA-seq expression values for *Hif1a*, *Hk1*, and *Hk3* in BMDMs treated with Lipid A for 4 h, with or without BRM014.

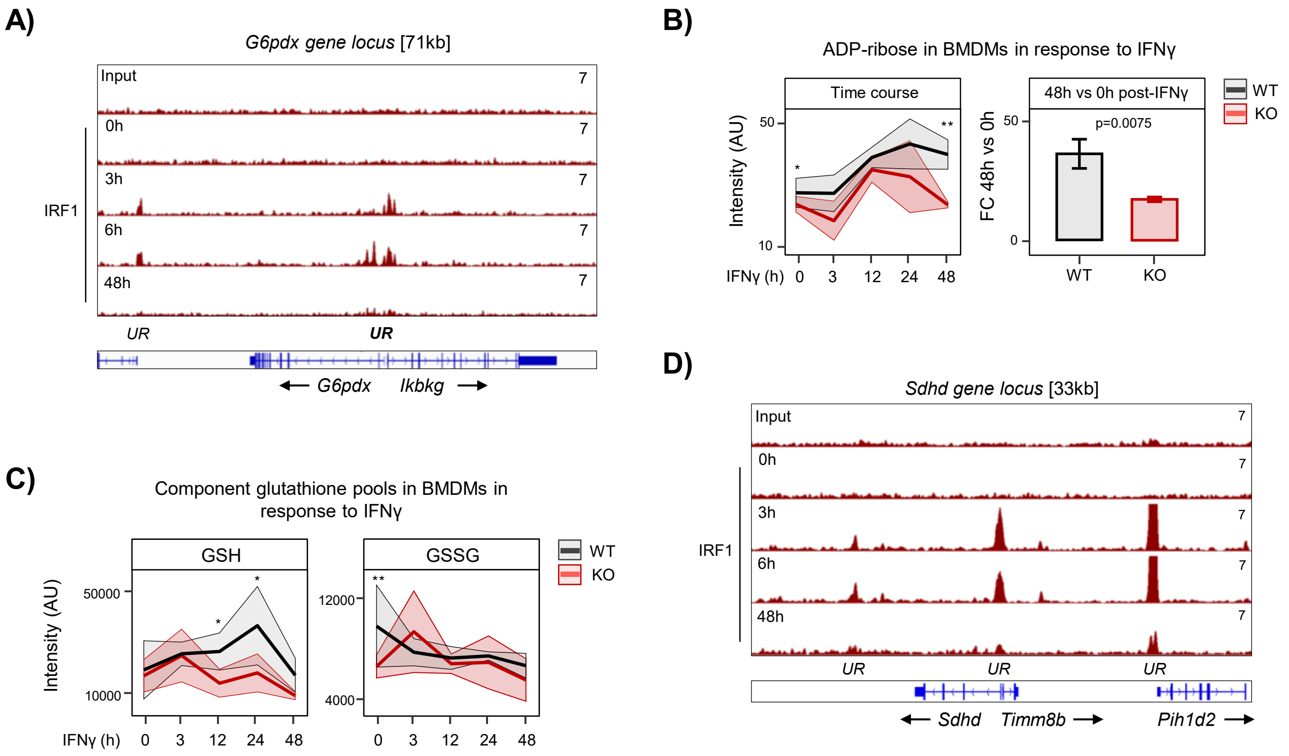


**Supplementary Figure 7. IRF1-dependent regulation of pentose phosphate, TCA, and redox cofactor metabolism, related to Figure 7.**
**(A)** Genome browser tracks showing IRF1 ChIP-seq signal at the *G6pdx* locus across IFNγ stimulation time points (0, 3, 6, 48 h) in BMDMs, with input control shown above. Gene body structures for *G6pdx* and *Ikbkg* and transcriptional orientation are indicated. [UR = Unresponsive] **(B)** Ribbon graph showing ADP-ribose levels in WT (gray) and IRF1 KO (red) BMDMs over 48 h following IFNγ stimulation, and bar graph quantification at 48 h. Error bars indicate SD. **(C)** Ribbon plots of reduced glutathione (GSH) and oxidized glutathione (GSSG) levels in WT and IRF1 KO BMDMs following IFNγ stimulation. Shaded regions represent standard deviation across biological replicates. [Metabolite intensity normalized by internal standard / median intensity; n= 3] **(D)** Genome browser tracks showing IRF1 ChIP-seq signal at the *Sdhd* locus following IFNγ stimulation across indicated time points. Gene body structures for *Sdhd*, *Timm8b*, and *Plp1d2* with transcriptional orientation are shown. [UR = Unresponsive] *$p < 0.05$, **$p < 0.01$.

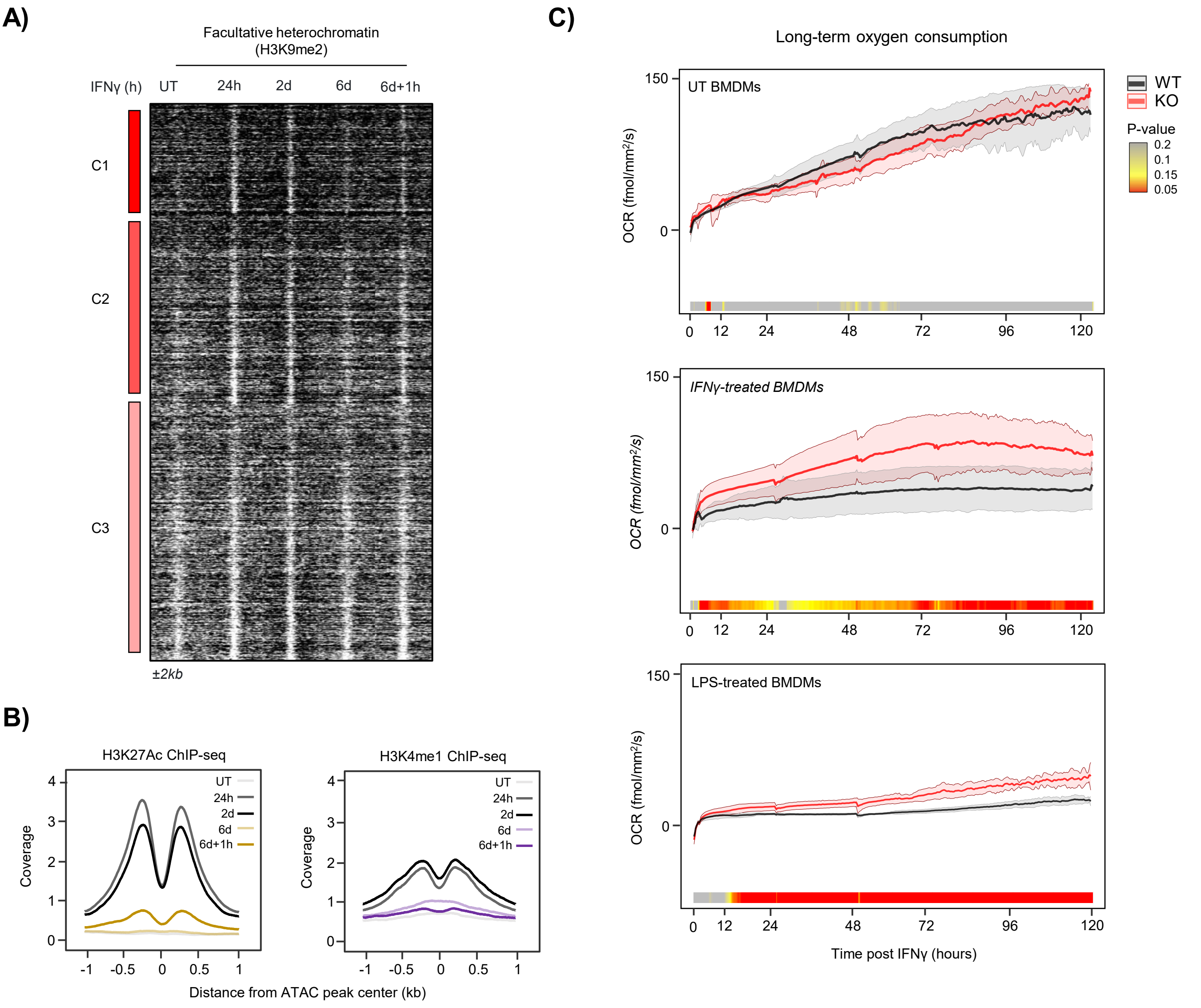


**Supplementary Figure 8. IRF1 sustains metabolic reprogramming and preserves active chromatin, suppressing facultative heterochromatin, related to Figure 8.**
**(A)** Heatmap of H3K9me2 signal across C1–C3 peaks during IFNγ stimulation and washout conditions. Signal intensity is shown across the indicated time points. **(B)** Aggregate ChIP-seq signal graphs of H3K27ac (left) and H3K4me1 (right) across indicated treatment and washout conditions. Traces correspond to untreated, stimulated, washout, and restimulation time points. **(C)** Ribbon graphs showing long-term oxygen consumption rate (OCR) kinetics (mean ± SEM, n = 4 per condition) for WT (grey) and IRF1 KO (red) BMDMs under untreated, IFNγ-activated, and LPS-activated conditions. Heat maps below each plot display rolling p-values from T-tests for WT versus KO comparisons, with red indicating $p < 0.05$.

| Target (gene or site) | Forward primer (5′–3′) | Reverse primer (5′–3′) | ChIP-qPCR for |
|---|---|---|---|
| ***Pomc*** | AGGCAGATGGACGCACATAGGTAA | TCCACTTAGAACTGGACAGAGGCT | IRF1, H3K4me1, H3K27ac |
| ***Tlr4*** | GTCAGCAAACGCCTTCTTCCTGTT | AGAGGAAGTGAGAGTGCCAACCTT | IRF1, H3K4me1 |
| ***Cd40*** | CTTCAGCTGTGGTCTTTCCCGTTT | ATCTCTGCAGAACCGAAAGCGTCT | IRF1, H3K4me1 |
| **Cluster 1 enhancer** | CAGGCCTAGGAGTCAGAGATATAG | CCATTGTTTGCAGGCATCAC | BRG1 |
| **Cluster 2 enhancer** | AGTCAAACTACTCCTGCACAT | AGTCAGGGACTCATCCTCAA | BRG1 |
| ***Clic5*** | GAGCACAGTATATCTGTATCATAGT | GAATTCCTCATTCTCCTATCTCAA | BRG1 |
| ***Nos2*** | AGCTGCTGGTACTGTGATTG | CTGCACTCCCAGACAAGTAAG | BRG1 |
| **4× IRF1 motif site** | CATGTGGACGTGAATACAAAGAATAAG | GCAATCACCCACCTGCTAAA | BRG1 |
| ***Tlr9*** | TTGTACCACCTGCTCTTTCAGGGT | AACTGGGCGGCAGAGAATGATGTT | H3K27ac |
| ***Cxcl9*** | TAGCTTGCAGAGGAAGTGCGGTAT | ATCCTTGGCTTTCTTCCCAGGTCT | H3K27ac |
| **Ubc (negative control)** | GTGTGACTGGACTCGGTAATAG | CTCCCTCTCTTTGCTCTACAAG | H3K9me2, H3K27me3 |
| **Tmem132b** | GTACACCCAAGGGAATCTACAC | GTGGACCACTGTGCCTATTT | H3K9me2 |
| ***Lrfn5*** | CAACACGATCTCTGGTGTATGA | CTAGCATTCGAGCCTGACTTAG | H3K9me2 |
| **H3K27me3 site 1** | TCCACGATTGGAAACCTCTAAC | CTAACGCATCCAACTTTCTTAACC | H3K27me3 |
| **H3K27me3 site 2** | CTGCCTCTGTACAATGGGATAA | TCCGACTCTCTGGGTCTTT | H3K27me3 |

**Supplementary Table 1. Primers used for ChIP-qPCR quality control and validation**
